# Scalable high-performance single cell data analysis with BPCells

**DOI:** 10.1101/2025.03.27.645853

**Authors:** Benjamin Parks, William Greenleaf

## Abstract

The growth of single-cell datasets to multi-million cell atlases has uncovered major scalability problems for single-cell analysis software. Here, we present BPCells, a package for high-performance single-cell analysis of RNA-seq and ATAC-seq datasets. BPCells uses disk-backed streaming compute algorithms to reduce memory requirements by nearly 70-fold compared to in-memory workflows with little to no loss of execution speed. BPCells also introduces high-performance compressed formats based on bitpacking compression for ATAC-seq fragment files and single-cell sparse matrices. These novel compression algorithms help to accelerate disk-backed analysis by reducing data transfer from disk, while providing the lowest computational overhead of all compression algorithms tested. Using BPCells, we perform normalization and PCA of a 44 million cell dataset on a laptop, demonstrating that BPCells makes working with the largest contemporary single-cell datasets feasible on modest hardware, while leaving headroom on servers for future datasets an order of magnitude larger.

## 1 Introduction

Improved single-cell genomics methods have increased dataset sizes by approximately two orders of magnitude in the last decade, but the scalability of analysis software has largely failed to keep pace. Today’s single-cell technologies routinely yield immense datasets, with contemporary single-cell atlases already exceeding 44 million cells [1]. In principle, these reference datasets should be broadly useful to and reusable by researchers around the world. In practice, however, memory limitations and other software scalability issues make even basic analysis with standard tools impossible on such large datasets.

At the time when popular single-cell analysis tools such as Seurat [2] and Scanpy [3] were designed, all but the largest single-cell datasets could comfortably fit in random access memory (RAM) on a laptop, making a primarily in-memory computational workflow logical for both speed and ease of implementation. However, as dataset sizes have grown to millions of cells, the extreme memory usage of these in-memory workflows has forced researchers to resort to lower-precision approximation methods or data subsetting to make analysis feasible on readily available hardware, partially negating the benefit of collecting large atlas datasets in the first place. Further improvements to single-cell data generation technologies promise still larger datasets [4, 5], creating a critical need for analysis software capable of scaling to handle the datasets of today and tomorrow.

To address this challenge, we created BPCells, a package for high-performance single-cell analysis of RNA-seq and ATAC-seq datasets. BPCells uses a two-pronged approach to provide fast, scalable single-cell analysis. First, BPCells uses a C++-based disk-backed streaming approach to carry out large-scale single-cell computations (including data normalization, PCA, and differential testing), while only loading a fraction of the dataset from disk into memory at a time. Second, BPCells introduces novel single-cell matrix and fragment storage formats based on bitpacking compression [6], which provide competitive space savings compared to other compression schemes with much faster read-/write speeds, further accelerating disk-backed computation. Given a fixed amount of RAM, the BPCells disk-backed approach increases the analyzable dataset size by nearly two orders of magnitude and increases the speed of marker feature tests, atlas-scale gene expression variance, and ATAC-seq peak matrix calculations by approximately 2x, 10x, and 50x respectively.

While disk-backed computations and novel storage formats can be used independently in the BPCells software package, unless otherwise noted our benchmarks utilize both innovations combined to showcase optimal performance. First, we benchmark our disk-backed streaming-compute implementation on single-cell compute tasks such as dataset normalization, PCA, and differential expression testing to compare BPCells with popular single-cell analysis tools across memory usage, speed, and scalability (2.1). Second, we analyze the bit-packing compression formats, measuring space savings, compression time, and decompression time relative to existing single-cell storage formats and general-purpose compression algorithms (2.2). Finally, we highlight how the combination of BPCells’s streaming computation and bitpacking compression enable a massive leap in scalability by demonstrating data processing, normalization, and PCA of the 44 million cell CELLxGENE census dataset on a mass-market laptop (2.3).

## 2 Results

### 2.1 Low-memory streaming compute

In-memory analysis tools, such as Scanpy or Seurat [2, 3], rapidly become memory-limited for contemporary large-scale datasets. For example, simply loading the 44 million cell scRNA-seq dataset [1] illustrated in Fig 1a into memory requires 750GB of RAM – massively outstripping the typical 16-32GB of RAM found on modern consumer hardware, and more than the 256GB available on the data center servers used in this paper’s benchmarks (see Methods). This high memory usage represents a key scalability bottleneck, and existing tools that try to reduce memory usage such as DelayedArray result in dramatically slower execution speeds – effectively trading one barrier to large-scale single cell data analysis for another.

**Fig. 1.**
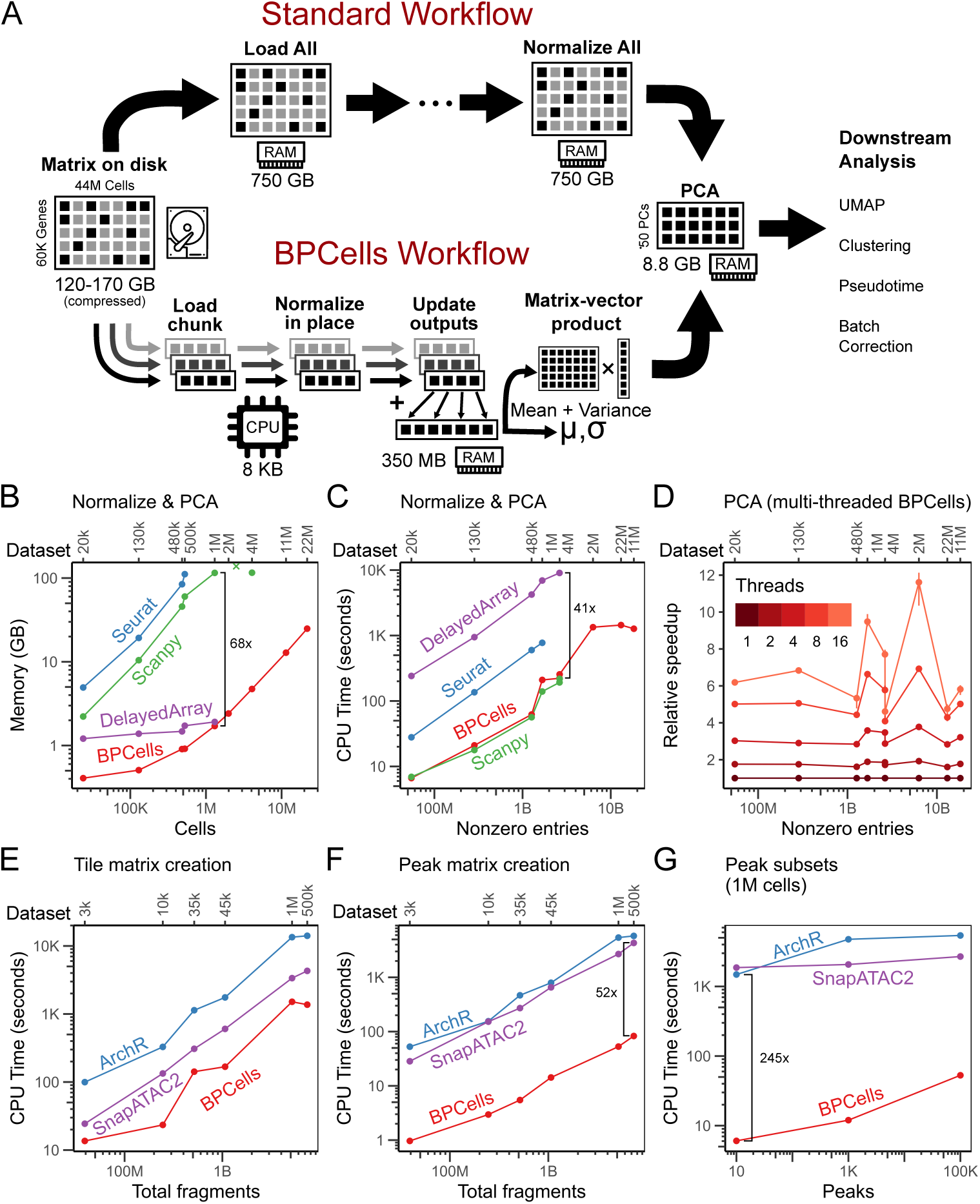
Streaming disk-backed computation eliminates memory bottlenecks while retaining fast execution speed. (a) Schematic of the BPCells streaming workflow compared to inmemory workflows. BPCells uses a streaming disk-backed workflow to eliminate the memory bottleneck between on-disk data and reduced dimensions. (b-c) Memory usage and time required for normalization and PCA of scRNA-seq data with various tools. BPCells shows (b) reduced memory usage, while (c) single-threaded execution speed remains comparable. Note differing x-axis scales since CPU time scales more directly with nonzero entries than cells. Seurat was evaluated with its in-memory workflow, without its optional disk-backed BPCells integration. (d) Speed-up in BPCells PCA calculation using multithreading. Multi-threading runs up to an order of magnitude faster than single-threading. (e) Tile overlap matrix and (f) peak overlap matrix creation time (100k peaks) across scATAC-seq tools. (g) Time required to calculate peak matrix subsets using different tools. BPCells’s fine-grained indexing enables even larger relative time savings for calculations on small subsets of the genome.

BPCells disk-backed streaming algorithms solve this challenge by providing a fast, low-memory way to perform computations on ATAC-seq fragments and single-cell feature×cell matrices. Each algorithm takes as input either a matrix or a fragment set in the form of a data stream of (feature, cell, value) or (chromosome, start, end, cell) data points. These data streams can either be produced from reading a data file on disk (see Table S7 for supported data formats), or taken from the output of another streaming algorithm (e.g. filtering and normalization operations, see Tables S3, S5 for supported matrix and fragment transformations). These operations can be assembled into an execution pipeline that passes data streams between each step, finally feeding into a stream-consuming operation such as differential feature testing, PCA, or saving transformed data to disk (see Tables S4, S6).

To put this in more concrete terms, consider an example normalization and PCA workflow illustrated in Fig 1a. First, counts for each cell are scaled to sum to 10k, then per-gene mean and variance is calculated to identify variable features. Next, the matrix is (optionally) subset to the most variable genes, log-transformed, then z-score normalized prior to running PCA. In-memory tools like Seurat or Scanpy perform all useful computations with in-memory, uncompressed sparse matrices, creating intermediate copies of the full matrix after every mathematical operation involved in matrix normalization.

In contrast, BPCells sequentially reads through the input file in chunks, passing each small chunk of data fully through its pipeline of streaming operations before loading the next chunk into memory. For example, to calculate mean expression per gene of a normalized scRNA-seq matrix, BPCells first loads a chunk of (feature, cell, raw count) data points, modifies the data in-place to get (feature, cell, normalized count), then updates a vector of per-feature sums based on the feature index and normalized count for each data point. After scanning through the entire matrix, this vector of per-feature sums can be rescaled to return per-feature means.

This streaming approach enables memory requirements to scale with the size of in-memory outputs requested by the user, rather than the total size of the input counts matrix or fragment file. With the exception of PCA, most operations can be completed with a single streaming pass over the dataset, and at most three inmemory tallies per cell (allowing analysis of about 40M cells per GB of RAM). PCA becomes the most expensive operation, taking approximately 250-300 passes over the dataset to compute 50 principal components with standard solvers, and requiring memory to store the principal components matrix for downstream analysis (about 5M cells per GB of RAM, assuming 50 principal components stored as 32-bit floats). As the size of a principal components matrix is ^-^85 times smaller than storing a feature×cell matrix in memory (Fig 1a), the BPCells streaming approach enables analysis of large-scale datasets by bypassing the memory bottlenecks caused when standard in-memory tools store a full data matrix in RAM (limiting them to at most ^-^60k cells per GB of RAM, assuming 2k features per cell). BPCells does not seek to address scalability in single cell analysis operations that can be performed directly from a principal components matrix or other reduced dimensions, such as clustering, UMAP, or pseudotime analysis (Fig 1a), as these operations do not represent the current scalability bottleneck given the small memory footprint of principal components matrices.

BPCells is able to preserve modularity without compromising performance by making all streaming algorithms read and write using C++-level data streams (see Supplemental Discussion). Users can freely recombine any of the supported operations (Tables S3-S6) to perform exploratory data analysis with a wide variety of normalization and data filtering strategies.

#### Normalization and PCA benchmark

We compared memory usage and execution speed of BPCells to existing analysis tools (Seurat, DelayedArray, and Scanpy [2, 3, 7])) on a range of nine scRNA-seq datasets ranging in size from 20k to 22M cells (Table S1). Each tool was tasked with performing a standardized normalization, variable gene selection and PCA workflow within a 3 hour time limit and 256GB memory limit (see Methods). We observed that BPCells’s memory usage scales with number of cells, whereas running time tends to scale with number of nonzero matrix entries (Fig 1b+c).

We evaluated Seurat’s in-memory workflow only, as disk-backed options in Seurat v5 use BPCells to perform normalization and PCA. Seurat’s in-memory workflow crashed on all datasets larger than 500K cells, as R’s Matrix package produces integer overflow errors for sparse matrices with over 2^31^ non-zero entries.

The DelayedArray package [7] offers disk-backed matrix operations in R that successfully reduce memory usage to a similar level as BPCells (Fig 1b), but at the cost of a substantial 41x slowdown relative to BPCells (Fig 1c). Despite using superficially similar disk-backed approaches, DelayedArray could not complete normalization and PCA of any dataset larger than the 1M neurons dataset within our 3 hour time limit, whereas BPCells was able to complete a 22M cell analysis in just 24 minutes (Fig 1c).

Scanpy was the best performing alternative to BPCells, with an execution speed about 17% faster on average than single-threaded BPCells (Fig 1c), but with 68x higher memory usage on a 1M neuron dataset (Fig 1b). Scanpy used ^-^5x more memory than the size of a single copy of the sparse data matrix, consistent with 5-10x excess memory requirements for the Python data analysis tool Pandas [8]. As a result, Scanpy could not analyze our 2M, 11M, or 22M cell datasets within the 256GB RAM limit of our benchmarking server.

Because BPCells memory usage scales with the size of the output principal components, BPCells can process a 22M cell dataset with about half the memory usage Seurat and Scanpy require on a 480K cell dataset (Fig 1b). Furthermore, BPCells achieves this performance improvement with no approximations or loss of precision, and only a modest increase in computation times compared to the fastest other tool we measured, when running with a single thread. BPCells allows additional speed increase on compute-intensive operations like PCA through the use of parallelization. On our benchmark datasets, we measured up to 12x faster PCA execution time when using 16 threads (Fig 1d), and just two threads were sufficient to make BPCells superior on both memory and execution speed for normalization and PCA across all tools and datasets tested. (See Supplemental Discussion for comments on file caching and multi-threaded performance.)

#### Randomized PCA with Scanpy/Dask

We also benchmarked an experimental integration of Scanpy with Dask [9], which aims to provide a disk-backed alternative to Scanpy’s default in-memory mode. Unfortunately, Scanpy version 1.10’s PCA demonstration with Dask [10] cannot perform full-precision PCA, instead using a randomized (rPCA) algorithm [11] with all accuracy-improving iterations skipped. This results in a poor approximation of the true principal components beyond the first dimension (Fig S1c). On our 1M-cell dataset which closely matches the size of the dataset used in the Scanpy with Dask demonstration, BPCells had 24x lower memory usage and 3.9x faster execution speed when running the same low-precision algorithm, and 26x lower memory and 1.8x faster speed when running a full-precision PCA (Fig S1a+b). This Scanpy with Dask setup was unable to complete the analysis of our 11M or 22M cell datasets within the 256GB of RAM available on our benchmarking server.

#### scATAC-seq matrix creation

For scATAC-seq analysis, the normalization and PCA steps are conceptually similar with a different set of common normalization choices. However, ATAC-seq analysis first requires generating the data matrix from fragment coordinates – tallying the number of genomic fragments overlapping regularly spaced genomic tiles (a “tile matrix”), or a set of high-signal peaks (a “peak matrix”). Here, we compared peak and tile matrix creation performance against ArchR [12] and SnapATAC2 [13], which use disk-backed methods for matrix creation from fragments, making speed, rather than memory-usage, the primary concern. (We omit dimensionality reduction comparisons, since SnapATAC2 and ArchR use in-memory approaches with similar memory bottlenecks to the previously-discussed RNA tools [13].) Because BPCells stores fragment data in genome-sorted order, it is able to perform an efficient linear scan over sorted fragment and peak/tile coordinates (see Supplemental Discussion for algorithm details).

We compared these matrix creation tasks across six scATAC-seq datasets ranging in size from 3k to 1M cells (Table S2). BPCells was 3x faster than SnapATAC2 and 10x faster than ArchR for tile matrix generation on our 500k cell dataset – the largest by total measured fragments (Fig 1e). For peak matrix calculation, BPCells was 52x and 70x faster than SnapATAC2 and ArchR, respectively (Fig 1f). As SnapATAC2 and ArchR use disk-backed techniques for peak and tile matrix creation, memory usage is sufficiently low for all tested tools that it is unlikely to become a bottleneck (Fig S1e+f).

The speed gap widens further when considering a smaller subset of the genome. To compute a 10-peak subset of the peak matrix on the 1M cell dataset, SnapATAC2 takes 31 minutes, ArchR 25 minutes, and BPCells 6 seconds (Fig 1g). This dramatic improvement is also enabled by storing fragments in fully genome-sorted order, which allows efficient lookup of all data overlapping selected genome regions without reading all fragment data. This allows BPCells to perform real-time interactive data visualization of large datasets directly from fragments stored on disk.

#### Marker feature testing

BPCells disk-backed streaming algorithms can also be used for statistical tests of marker and differential features, such as Wilcoxon rank sum tests. BPCells Wilcoxon calculations use a similar algorithm to Presto [14], performing a sparsity-aware rank transform for each gene followed by a one-vs-rest statistical test for over- and under-expressed features in each cell type. On our 500k cell benchmark dataset, Presto and Scanpy used 154x and 80x more memory, respectively, than BPCells, while taking 2.3x and 40x more time, respectively (Fig S1d). This speed advantage of BPCells relative to Presto is surprising due to the similarity in the algorithmic approaches. While we reserve full discussion of the likely reasons for this performance gap to Supplemental Discussion, two contributing factors are BPCells’s choice of sorting algorithm and support for input matrices in feature-major storage order (as opposed to cell-major; see “Data storage ordering” below and Supplemental Discussion).

### 2.2 High-speed bitpacking compression

For disk-backed workflows, disk read/write speeds become a likely bottleneck that can cause slower execution speeds. When running CPU-light pipelines, a single BPCells thread can process over 3 GB/s of uncompressed matrix data, 40% of the 7 GB/s rated sequential read bandwidth of the laptop SSD storage used for some benchmarks in this study (see Methods). With the addition of multi-threading, the compute throughput can rise over an order of magnitude, making the fixed storage bandwidth a serious potential bottleneck to execution speeds. Compression algorithms can mitigate this bottleneck by trading some compute capacity to reduce the storage size of files on disk and hence reduce the required read bandwidth for disk-backed workflows. Unfortunately, general-purpose compression algorithms like gzip [15] tend to require too large a compute cost to make the storage bandwidth reductions practically useful for a tool like BPCells.

Since general-purpose compression algorithms are well-studied and well-optimized, we opted to develop a specialized compression algorithm that could take advantage of known properties of single cell data types to achieve faster compression and decompression. We built on top of the BP-128 compression scheme, a form of bitpacking compression introduced by Lemire and Boytsov [6] that is specialized to compress lists of 32-bit integers, providing better space savings for inputs that are lists of small positive integers (or easily transformed into small positive integers). BP-128 can be implemented with SIMD CPU instructions to achieve exceptional compute efficiency of fewer than 2 CPU cycles per integer [6], and its small block size of 512 bytes enables efficient random access for small reads if necessary. We identified practical transformations that enable single-cell datasets to be compressed via BP-128, with BPCells’s implementation achieving comparable space savings to general-purpose algorithms at dramatically higher compute efficiencies.

#### Bitpacking compression of matrices

The BPCells bitpacking strategy for matrices (Fig 2a) starts with a standard compressed sparse column (CSC) layout, similar to that used by the AnnData h5ad and 10x HDF5 matrix formats. Two integer arrays store the row index and value for each nonzero entry in the matrix, with the entries ordered by column with rows as tiebreaker. A third, shorter “column start” array stores the starting index of each column within the row and value arrays (Fig 2a). This sparse matrix layout avoids wasting storage space on zero-valued entries, as typically fewer than 7% of values in a single cell counts matrix are non-zero (Table S1). BPCells optionally stores row and column names to help track feature and cell IDs, but only the row and value arrays are compressed as all other elements typically make up a negligible fraction of the total data size.

**Fig. 2.**
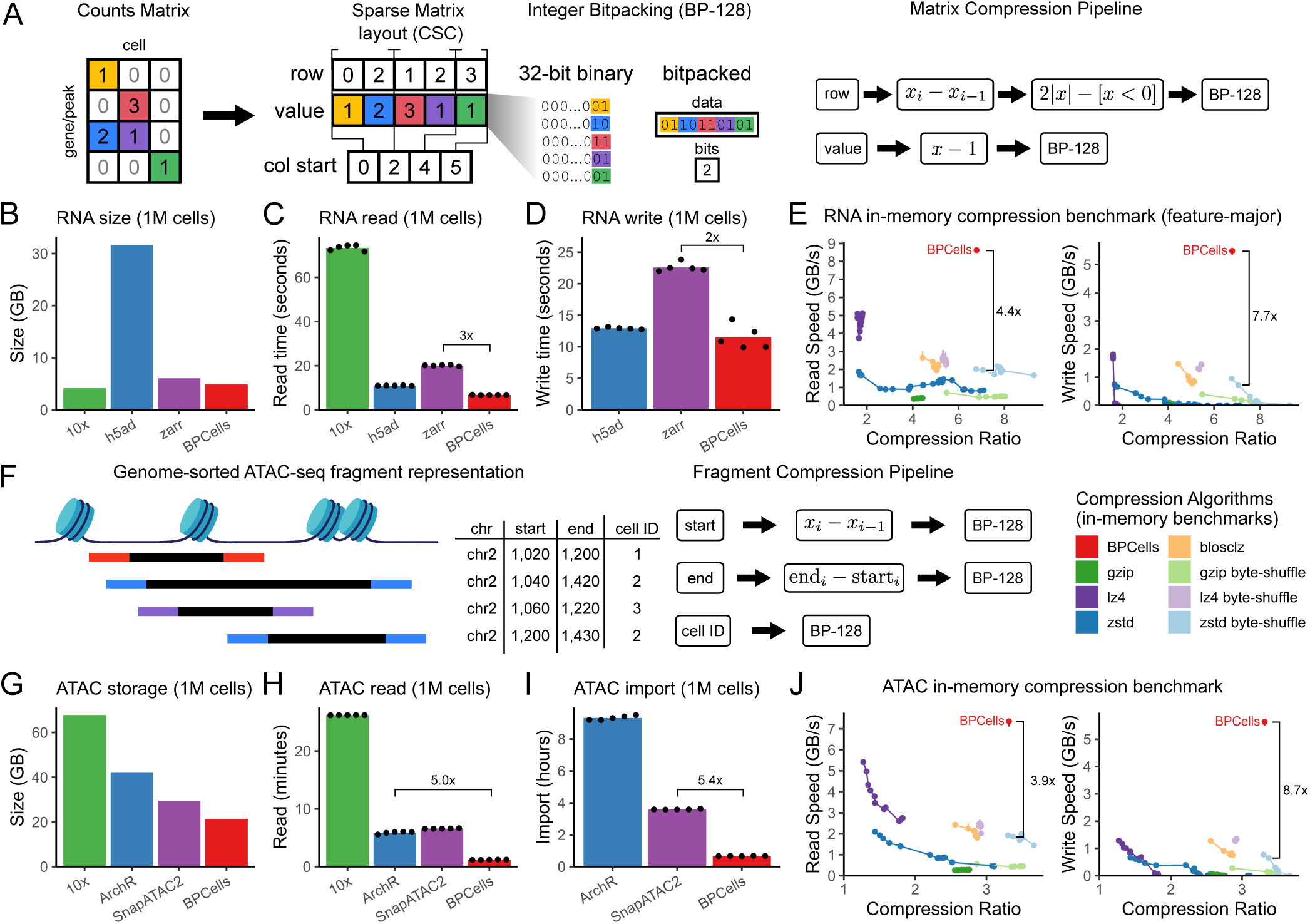
Bitpacking compression allows high-speed and compact single-cell file storage. (a) Bitpacking schematic overview. BPCells bitpacked matrices begin with a standard compressed sparse column layout. Then each array is bitpacked to remove leading 0-bits across 128-integer chunks and the number of bits used is separately recorded. A pipeline of transformations improves compressibility of the row and value arrays prior to bitpacking. (b-d) File format performance on a 1M cell scRNA-seq matrix for (b) size, (c) read speed, and (d) write speed. Comparison single-cell file formats used with default compression settings. (e) Compression algorithm compute-efficiency benchmark on a 130k cell scRNA-seq matrix stored in feature-major order. Read/write speeds measured in uncompressed bytes, and compression ratio as uncompressed size/compressed size. Dots indicate distinct compression level settings when available. (f) BPCells bitpacked scATAC-seq fragments store coordinates in genome-sorted order, followed by a transformation pipeline and bitpacking in 128-integer chunks. (g-i) File format performance on a 1M cell scATAC-seq dataset with (g) size, (h) read speed, and (i) write speed. (j) Compression algorithm compute-efficiency benchmark on a 10k cell scATAC-seq dataset. Read/write speeds measured in uncompressed bytes, and compression ratio as uncompressed size/compressed size. Dots indicate distinct compression level settings when available.

To perform bitpacking with BP-128, BPCells first calculates how many bits are required to represent the largest value in a chunk of 128 integers. It then discards the many leading zero bits from the native 32-bit representation (“packing” the variable bits to be stored contiguously), while separately storing the bit width for use in later decompression (Fig 2a). BPCells implements separate data transformations for the row and value arrays to improve space savings, which are outlined in Fig 2a and described in detail in Supplemental Discussion. In brief, these transformations result in the value array compressing best for small, non-zero counts values and the index array compressing best when consecutive entries have small positive differences (Fig S2e+f). For matrices holding non-integer data, the value array is left uncompressed.

With the BPCells matrix compression format, we observe space savings of approximately 4x for our benchmark RNA-seq datasets given the original cell-major data ordering, but this can be improved to a median of 6-7x space savings by either transposing to a feature-major ordering or sorting the rows (i.e. the non-major axis) by the mean value within each row (Fig S2a+b). On scATAC-seq peak and tile matrices, we achieve 4x to 6x space savings with the BPCells compression scheme, and more modest savings from switching away from a cell-major storage order (Fig S2c+d). The space savings from bitpacking scale consistently with simple summary statistics across scRNA-seq and scATAC-seq counts matrices (Fig S2e+f), suggesting that the BPCells compression scheme is not overly specialized to RNA-seq and ATAC-seq matrices and should achieve predictable performance on generic sparse integer matrices.

We compared the size and speed of BPCells compressed matrices to the 10x HDF5, AnnData h5ad, and AnnData Zarr formats according to each format’s default compression settings (10x: gzip, h5ad: uncompressed, Zarr: Blosc-LZ4). On a 1 million cell dataset, BPCells used 4.9GB of storage, second only to the 10x HDF5 format at 4.2GB (Fig 2b). However, the 10x format exhibited the slowest read speed by far at 73 seconds, with Zarr at 20 seconds, uncompressed h5ad at 11 seconds and BPCells at just 6.8 seconds (Fig 2c). For writing, Zarr took 23 seconds, uncompressed h5ad 13 seconds, and BPCells 12 seconds (Fig 2d). The 10x format was omitted from write benchmarks due to lack of a first-party tool for isolated matrix write operations. These results show that the combination of good space savings and extreme compute efficiency for bitpacking allow BPCells to eliminate the speed-vs-size trade-off present in the other compressed formats – BPCells saves both disk space and time reading/writing from disk.

#### Bitpacking compression of scATAC-seq fragments

BPCells fragment set storage consists primarily of three integer arrays containing genomic start and end coordinates, and a numeric cell ID for each fragment (Fig 2f). The fragments are stored in genome-sorted order by start coordinate, allowing chromosome data to be represented with a short array listing the storage range of each chromosome in the start, end, and cell arrays. To allow efficient subsetting by genomic region, BPCells also includes an end-coordinate indexing array 1*/*128^th^ of the length of the main start, end, or cell data arrays. Textual cell names (often barcode sequences) are stored separately to allow conversion with the numeric cell IDs. See Supplemental Discussion for format details.

Before performing BP-128 compression, BPCells adds data transformations to the start and end data arrays (Fig 2f, Supplemental Discussion). With these transformations, the start coordinate array compresses best when the distance between consecutive start coordinates in the sorted list is small, as is common at high-accessibility sites in scATAC-seq data (Fig S2g). The end coordinate array compresses best when fragment lengths are small (typically *<*1kbp for ATAC-seq [16]), and the cell array compresses best at lower cell counts (Fig S2h). Overall storage per fragment is driven mainly by mean fragments per cell, such that filtering out fragments from low-read, non-cell barcodes modestly reduces per-fragment storage size by ^-^18% (Fig S2i).

The BPCells fragment file format differs notably from ArchR arrow files and SnapATAC2, which group fragments by cell ID rather than globally sorting by genome location. This cell-based grouping can virtually eliminate the per-fragment storage costs of cell IDs, but it causes a major slowdown when performing reads of small genome subsets (Fig 1g). With the bitpacking compression used in BPCells, we observe that improved compression of the start coordinate array almost perfectly compensates for increased cell ID storage costs caused by intermixing the fragments from higher numbers of cells (Fig S2h), leading to smaller total file size while maintaining the advantages of fully genome-sorted fragment storage.

We compared the performance of the BPCells fragment file format to 10x fragment files (gzipped text), ArchR arrow files (uncompressed HDF5), and Snap-ATAC2 (HDF5 with gzip). On a 1 million cell scATAC-seq dataset, we see that the BPCells compressed fragment file has the smallest storage size at 21GB (28% smaller than the next-closest method), has 5x faster read times than the next-closest alternative at 1.2 minutes and 5.4x faster import times than the next closest alternative at 40 minutes (Fig 2g-i).

#### Comparison with general-purpose compression algorithms

To better understand the compute efficiency vs. space savings trade-off of bitpacking compression, we ran inmemory compression benchmarks of BPCells compared to seven general-purpose compression algorithms with a total of 226 different configurations (82 of which are shown in Fig 2e+j, see Methods). The primary algorithms compared were gzip [15], LZ4 [17], and Zstd [18]. We also included Blosc variants of these core algorithms, which apply specialized compression-enhancing preprocessing filters prior to running a general-purpose compression algorithm [19]. Each non-BPCells algorithm included a compression level parameter, which we varied to explore different trade-offs between space savings and compute efficiency.

BPCells offered the best read and write speeds across all tested compression algorithms, and competitive space savings compared to the best algorithm variants. On the three tested storage examples, BPCells decompression produced data at 7.4-8.6 GB/s and BPCells compression consumed data at 5.1-5.6 GB/s (Fig 2e+j, S3a).

Although other algorithms can produce better space savings compared to BPCells at their highest compression settings, this comes at the cost of dramatically slower write speeds. Zstd compression level 9 with Blosc byte-shuffle filter provided the best space reduction at 27% and 9.6% smaller than BPCells on feature-major matrices and scATAC-seq fragments, but at the cost of 102,500% and 68,900% slower writes (Fig 2e+j). For cell-major matrices, Zstd compression level 22 provided the best space savings at 42% smaller than BPCells, but 304,500% slower writes (Fig S3a).

Overall, these in-memory benchmarks show that BPCells outperforms sophisticated general-purpose compression algorithms with a simple, high-performance bitpacking algorithm specialized to single-cell data types.

#### Data storage ordering

While most BPCells streaming algorithms are order-independent and agnostic to the order data points are processed, certain algorithms require particular data orderings to allow efficient execution as previously highlighted (Figs 1f-g+S1d, Tables S3+S4). As BPCells mainly performs sequential scans of files on disk, establishing the correct storage order for order-dependent algorithms such as marker feature testing requires fast, low-memory algorithms for sorting and re-ordering data.

For matrices, BPCells offers two storage orders: the column-major compressed sparse column (CSC) layout illustrated in Fig 2a, and the analogous row-major compressed sparse row (CSR) layout. In the column-major layout, each column of matrix entries is stored contiguously, meaning a single column of the matrix can be loaded quickly with a small sequential read, but loading a single row of the matrix is no faster than reading the full matrix. Row-major layouts have the inverse properties, with fast row reads and slow column reads. Because R and Python differ in convention of whether cells correspond to matrix columns or rows, we avoid ambiguity by referring to cell-major and feature-major layouts depending on whether cells or features respectively can be loaded efficiently.

There is no universal answer to whether cell- or feature-major matrix storage orders are best, so BPCells must provide an efficient way to convert between matrix storage orders. Seurat, Scanpy, AnnData, and 10x matrix files most commonly default to a cell-major order, and we have shown that a feature-major storage order can have advantages for compression efficiency and enabling disk-backed algorithms that are not order independent (notably Wilcoxon marker feature testing, see Tables S3+S4, Fig S2a). While transposing the storage order between row- and column-major is fast with in-memory algorithms, BPCells cannot rely on such in-memory approaches without introducing a major memory scalability bottleneck. To that end, BPCells implements a disk-backed storage order transpose operation which requires 147x less memory than SciPy [20] on a 2M cell dataset and uses a constant 1.4GB of RAM to sort datasets up to 22M cells (30GB compressed file size), while using just 39% and 52% more time, respectively, than SciPy and R dgCMatrix [21] in-memory transpose routines (measured by geometric mean, Fig S3b, see Supplemental Discussion for algorithm details). For fragment files, BPCells has a similar problem when allowing fragment files from many samples to be merged either on-the-fly or to create a single whole-dataset fragment file. The BPCells disk-backed fragment operations such as peak and tile matrix creation require genome-sorted input streams, so working with multi-sample datasets also requires an efficient way to merge fragment sets while maintaining genome-sorted order. We compared performance of the GNU sort [22] program to BPCells merging multi-sample datasets into a whole-dataset fragment file (see Methods). We observed approximately constant memory usage for both programs, but BPCells ran a geometric mean of 38x faster (Fig S3c, see Supplemental Discussion for algorithm details). These efficient storage reordering operations help ensure that BPCells has a complete toolkit for merging, filtering, and converting matrix or fragment files – a necessary complement to its disk-backed compute capabilities (Tables S3-S6).

### 2.3 Scalability to massive cell atlases

To assess the scalability of BPCells at the limits of contemporary single-cell datasets, we turned to the CELLxGENE census human dataset [1], a unified collection of over 44 million unique cells collected from hundreds of scRNA-seq datasets deposited with the Chan Zuckerberg Initiative (CZI). As analysis of this dataset is easily within the capabilities of BPCells on our benchmarking server hardware (256GB RAM), we also ran the workflow on a consumer laptop (32GB RAM) to allow us to test scalability limits. All benchmarks were run with parallelization enabled to achieve maximal performance.

#### Bulk matrix read and write

For our first series of benchmarking tasks, we performed bulk matrix read/write operations – first subsetting the census dataset to remove duplicate cells as a write benchmark, then measuring the size and read speed of the unique cells data matrix. We compared the performance of the BPCells format with the TileDB-SOMA format [23] developed for the CELLxGENE census as a collaboration between CZI and TileDB, Inc. While the TileDB-SOMA format was initially designed around network-based access, its incorporation of the widely used TileDB array storage system [24] allows it to serve as an optimized baseline for storage and querying of large sparse matrices from a fast local file-system.

For storage size, BPCells and the CELLxGENE census take different approaches but use roughly comparable storage space for the matrix of unique cells. The census TileDB format stores two copies of the data matrix: raw counts and normalized counts, which use 147GB and 166GB of storage respectively (Fig 3a). BPCells only explicitly stores the counts matrix, but provides two possible storage layouts: cell-major (168GB) and feature-major (117GB; Fig 3a). Both BPCells matrices are partitioned into 100k cell chunks to allow easier parallelization, while TileDB uses 2,048×2,048 tile sizes (see Methods).

**Fig. 3.**
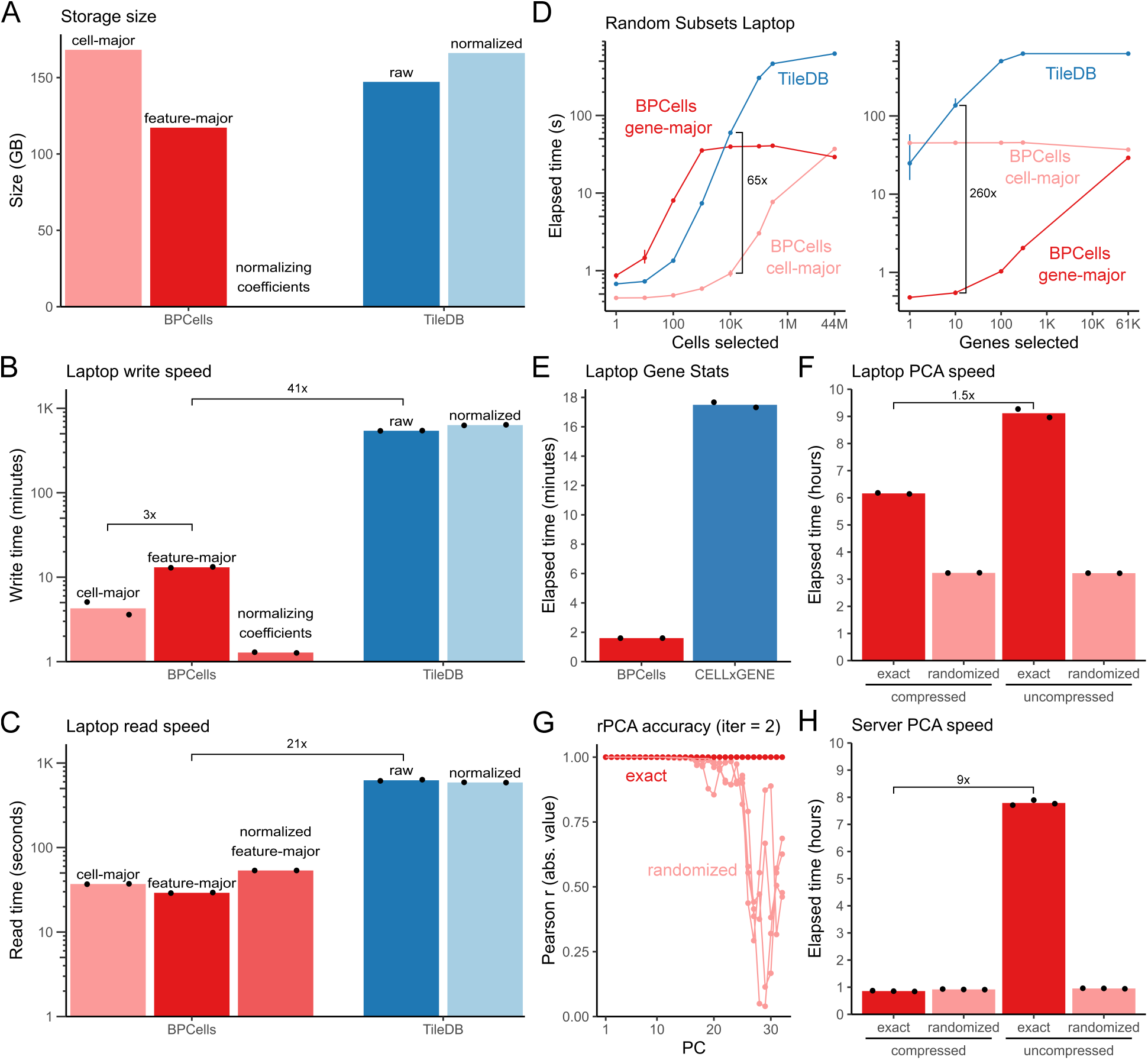
Performance of 44M cell analysis of CELLxGENE census on a laptop. (a-c) Performance of BPCells sparse matrix formats and storage layouts compared to TileDB, comparing (a) size, (b) write speed and (c) read speed (fully sequential) on a laptop. Write speed for feature-major includes time to swap storage order from cell-major to feature-major. (d) Read speed for random subset queries of cells and genes on a laptop. (e) Speed to calculate per-gene mean and variance on a laptop. (f) PCA on a laptop runs faster with compressed input, while randomized PCA (iter=2) is faster and less sensitive to input format. (g) Randomized PCA (iter=2) accuracy is high for early components but diminishes for later components. (h) PCA on a server runs faster than on a laptop, with a large advantage for exact PCA with compressed input compared to uncompressed input.

For our write benchmark task of subsetting to remove duplicate cells and writing a new matrix copy, we found dramatically faster write and read speeds for BPCells compared to TileDB. BPCells offered speedups of 41-125x and 11-21x for writes and reads respectively on a laptop (Fig 3b+c), and on our servers offered 63-186x and 15-41x speedups for writes and reads respectively (Fig S4a+b). Writing to BPCells feature-major layouts was modestly slower due to introducing a storage order transpose operation from our cell-major inputs. For reading, feature-major reads were modestly faster than cell-major reads at 1.3x or 2.6x faster on a laptop or server respectively (Fig 3c, S4b).

Rather than explicitly store normalized counts data, BPCells instead stores just the required normalization coefficients (i.e. per-cell total read counts), which take negligible storage space (0.35GB, Fig 3a), enabling the BPCells streaming architecture to quickly re-calculate full-precision normalized values on-the-fly when reading from the raw counts matrices. The time to compute and save these normalized values is comparable to the time to write cell-major subsets (Fig 3b, S4a), and data read takes 1.8-2.7x longer to calculate the normalization on-the-fly – still 11-15x faster than reading the pre-computed normalized TileDB matrix (Fig 3c, S4b). Thus, the streaming approach allows inclusion of normalized data at little-to-no cost, and the underlying counts matrix storage need only be duplicated if required for downstream query performance.

#### Matrix subsetting

Because one core intended application of the TileDB-SOMA storage layout is reading small subsets of the data matrix, we performed a direct comparison with BPCells on this task. We selected two types of sub-setting: subsetting along the cell axis only (e.g. as a mini-batch for training a neural network expression model), and subsetting along the gene axis only (e.g. to quantify per-cell expression of key marker genes). When querying by cell, the BPCells cell-major format can load a random 10,000 cell subset in 1 second on average, but TileDB averages 60 and 18 seconds on a laptop or server respectively (Fig 3d, S4e). The BPCells feature-major format can load 10 random genes in 0.5-0.7 seconds, whereas TileDB takes 143 and 54 seconds on a laptop or server respectively (Fig 3d, S4e). We also see performance advantages on sequential, rather than random, data subsets (Fig S4f+g). See Supplemental Discussion for recommended BPCells storage layouts for different applications.

#### PCA and matrix computations

We next performed normalization and PCA on the 44 million cell dataset, which allowed us to evaluate the performance of combining BPCells streaming computation and compressed formats with both modest compute resources (a consumer laptop) and more powerful server hardware. As an example of normalization performance, we calculated per-gene mean and variance across all 44 million cells in the census dataset, a common sub-task both for variable gene selection and for final z-score normalization prior to PCA. We observed BPCells running times of 96 seconds on a laptop and 32 seconds on a server for gene mean and variance (Fig 3e, S4c). As TileDB does not provide direct compute functionality, we compared to the CELLxGENE census Python package. BPCells used 4.6x and 5.8x less CPU time on a laptop or server respectively, with a larger gap in wall clock time (11x and 38x respectively) since CELLx-GENE currently only parallelizes data loading without parallelizing the mean and variance calculations (Fig 3e, S4c+d).

To demonstrate PCA calculations on the 44 million cell census dataset, we used standard log normalization and retained all ^-^60k genes rather than subsetting to 2,000 variable genes as in our comparison to in-memory tools. We reasoned that the biological diversity in all-tissue atlas datasets may not always be captured by ^-^3% of genes, and wanted to demonstrate the unique scalability of BPCells in this computationally-intensive all-gene PCA workflow. We computed the top 32 principal components in 6.2 hours on a laptop and 51 minutes on a server when using a compressed bitpacked feature-major input matrix (Fig 3f+h). Using a randomized PCA algorithm lowers compute time by reducing the number of required matrix passes from 167 to 6, allowing substantial time savings on a laptop at the cost of poorer accuracy, particularly beyond component 16 (Fig 3f-h).

For exact PCA calculations, we observed that using the BPCells compressed storage format provided end-to-end speedups of 1.5x and 9x on a laptop or server, respectively, compared to using uncompressed data storage (Fig 3f+h). This shows how BPCells high-performance compressed storage can speed up multi-threaded disk-backed streaming algorithms by reducing disk read bandwidth bottlenecks (see Supplemental Discussion).

Extrapolating from our benchmarks on smaller datasets, we estimate this 44M-cell PCA calculation would require over 4TB of RAM with in-memory Scanpy, and 1.5 to 10 days of time on a server or laptop respectively with DelayedArray. Additionally, none of the other tested tools were able to successfully complete full-precision normalization and PCA on our 2M, 11M, or 22M cell datasets within the benchmark server’s memory and time limits. Therefore, we conclude that BPCells is uniquely capable of scaling to our 44M cell analysis demonstration, and it can even achieve this level of scalability on a consumer laptop.

## 3 Discussion

BPCells aims to improve scalability across the single-cell software ecosystem by providing a modular interface to scalable, disk-backed streaming computations. For example, Seurat v5 [25] uses BPCells to provide disk-backed analysis as an alternative to many of their existing in-memory workflows. This promising early adoption of BPCells demonstrates its utility not only as a standalone single-cell analysis tool, but also as a scalable back end for existing analysis software. Because BPCells can directly read and write h5ad and 10x HDF5 matrix files, it is easy to get started analyzing existing data files as-is or to quickly convert into the higher performance BPCells formats.

Based on performance in the benchmarks presented here, we expect that BPCells will enable a wide variety of additional matrix normalizations and dimensionality reduction methods in a scalable, efficient manner, often by recombining the existing set of matrix and ATAC-seq fragment operations without changes to the core BPCells package. For example, non-negative matrix factorization [26], latent semantic indexing [27], and SnapATAC2’s cosine similarity-based dimensionality reduction [13] can be implemented in a disk-backed manner using existing BPCells operations (Table S3, S4). Minor extensions would enable the training of robust, atlas-scale cell type classifiers via generalized linear models [28] or correlating chromatin accessibility with gene expression across multiomic atlas datasets. The BPCells matrix and fragment-set operations can likely be used for additional single-cell modalities, such as scCUT&Tag or DNA methylation, and potentially even fields beyond single cell analysis that utilize large sparse matrix computations.

Efficient on-the-fly data normalization with BPCells streaming algorithms also allows for straightforward improvements to single-cell matrix storage formats that currently hold both raw counts and normalized data copies. Storing just the normalization formula and parameters rather than a full normalized matrix copy provides at least 2x space savings with minimal run-time cost for on-the-fly normalization. Furthermore, integer counts matrices are both more compressible than normalized matrices, and in multi-threaded workloads, on-the-fly normalization of bitpacked counts data is faster than simply loading uncompressed, normalized data due to disk bandwidth limitations. Instead of storing raw and normalized data, we suggest storing the raw counts data in both cell-major and feature-major storage orders, allowing downstream applications to optimize performance by choosing the best storage ordering based on the user’s requested operation.

We have demonstrated that BPCells can allow well-studied existing analysis methods such as principal components analysis to scale to dramatically larger datasets. In many cases, this capacity to calculate a full-precision, full-matrix PCA reduces or eliminates the need for sketch-based or other approximation methods for dimensionality reduction [25, 29]. Single-cell neural network methods [30] may also benefit from fast random access to single-cell data profiles during training using BPCells compressed data formats. We anticipate that the efficient BPCells storage and disk-backed compute infrastructure will help the single-cell computational community to develop new scalable analysis methods in the future. Overall, BPCells makes rapid, atlas-scale analysis accessible even to researchers with modest compute resources, expanding the reach and utility of massive public reference datasets.

## 4 Methods

### Benchmarking hardware

Benchmarks in this paper were run primarily on server hardware featuring AMD Zen 2 (Rome) EPYC 7502 CPUs with simultaneous multi-threading disabled, 256GB of RAM (8×32GB of registered 3200MT/s DDR4 with 1 DIMM per channel), and 2TB of local NVMe SSD storage (2x PCIe Gen3×4 drives in RAID 0). CELLxGENE census benchmarks were additionally run on a consumer laptop with an Intel i5-1240P CPU, 32GB of RAM (2×16GB of 3200MT/s DDR4), and 2TB of local NVMe SSD storage (PCIe Gen4×4, Samsung 990 Pro)

### Software versions

All benchmarks were performed on BPCells v0.3.0 and all other software library versions were taken from a container snapshot built in December 2024. For exact library versions and a runnable container file with the benchmarking environment, see “Data and code availability” below.

### Memory and running time measurement

Memory usage was measured at the process level using Max RSS from GNU time -v. Running time for specific code blocks in R was measured using system.time(), counting the sum of user and system time for CPU time and taking elapsed time as given. Running time in Python was measured using time.process time() for CPU time and time.time() for elapsed time.

### Dataset preparation

Dataset download and preparation code available under the datasets supplemental code directory. RNA counts matrices were downloaded from the original published source (Table S1), then converted into 10x HDF5 and BPCells-format matrix files. Initial variable gene selection was performed with Seurat’s BPCells integration (NormalizeData(), then FindVariableFeatures() with default parameters) to obtain variable features for the standardized PCA benchmark. Basic unsupervised clustering was performed with BPCells to obtain cluster assignments for marker feature benchmarking.

ATAC datasets were downloaded and converted to 10x- format and BPCells-format fragment files (Table S2). Fragment files were then filtered to barcodes passing quality control, from provided cell calls (3k PBMC, 10k PBMC, 1M brain), or via TSS enrichment and read count cutoffs (35k hematopoiesis, ENCODE 45k pancreas and 500k heart; see find-passing-cells.R in supplemental code). After filtering, samples from each dataset were separately converted into ArchR arrow and SnapATAC2 h5ad formats. Peak sets for each dataset were downloaded in BED format for peak matrix creation benchmarking.

### RNA Normalization & PCA benchmark

A standardized normalization and PCA workflow was performed with all tools, consisting of (1) Subsetting to genes with at least 1 detected read across all cells, (2) Scaling counts per cell to sum to 10k, (3) Calculating per-gene mean & variance, then subsetting to the pre-calculated variable gene set, (4) log1p-transforming the matrix, (5) Normalizing values for each gene to have variance of 1 and mean of 0, and (6) Performing PCA with 50 components. All tools were limited to 3 hours of execution time and 256GB of RAM.

When available, mean-centering was performed implicitly during the PCA step to improve performance (all tools except for Seurat). For BPCells and DelayedArray, a normalized copy of the matrix was saved to disk prior to mean-centering to avoid expensive re-calculation of log1p during each PCA pass. Performance measurements for additional BPCells variants with an intermediate variable-gene-subset counts matrix saved to disk or no intermediate matrix saved to disk are available in supplemental data (see Supplemental Discussion for details).

For the Scanpy with Dask benchmark, configuration parameters were modeled off of the “Using dask with Scanpy” tutorial in the Scanpy documentation. Code available in supplemental code rna-timing/pca-benchmark.

### Wilcoxon marker genes benchmark

Marker genes were calculated for a set of pre-calculated unsupervised clustering assignments on each dataset, using a gene expression counts matrix normalized so each cell adds to 1. Tools were limited to 3 hours of running time and up to 256GB of RAM. Scanpy and Presto used uncompressed 10x-format matrix input, while BPCells used the compressed BPCells file format as input in feature-major order. As Scanpy performance is dramatically lower when applying the standard statistical correction for ties, it was benchmarked both with and without tie correction. BPCells and Presto always perform tie correction. Code available in supplemental code rna-timing/marker-genes.

### ATAC peak and tile matrix benchmark

For each dataset, filtered fragment files in the appropriate format for each tool were used as input. Each sample from within the dataset was processed and timed independently, then CPU times were summed together per dataset for benchmark analysis. Each tool was configured to use a single thread for each sample. Tile matrices were computed with 500bp bins, and peak matrices were calculated for peak sets consisting of 100k, 1k, and 10 peaks sampled uniformly at random from the per-dataset peak sets. Code available in supplemental code atac-timing/peak-tile-timing.

### RNA 1M cell file format comparison

The 1M cell scRNA-seq data matrix was downloaded from 10x with original compression settings, then converted to uncompressed h5ad, LZ4-Blosc compressed Zarr, and bitpacking compressed BPCells matrices. Read times for 10x and h5ad formats were measured with both BPCells and ScanPy/AnnData respectively, taking the minimum observed running time across tools. BPCells read times were calculated as the time to compute column sums on the matrix values, and Scanpy/AnnData read times were calculated as time to load a matrix copy into memory. Write performance was measured as time to write in-memory matrix copies to disk. All times reported as CPU time; elapsed time statistics available in supplemental data. Code available in supplemental code compression/rna-1M-cell.

### ATAC 1M cell file format comparison

Unfiltered fragment files were first imported and subset to fragments falling on chr1-chr22, chrX, or chrY with no subsetting of cells by quality control thresholds. Read time was measured as time to load fragment data from disk into memory for BPCells, ArchR, and SnapATAC2, and time to run gunzip | wc −l for 10x. Each sample was run independently and total times for the 1M cell dataset were taken as the sum across all samples. All tools were configured to run single-threaded. We observed SnapATAC2 used multiple threads even when n jobs=1, so we report CPU time rather than elapsed time. Benchmarking results for additional datasets are available in supplemental data. Code available in supplemental code compression/fragments-read-write.

### In-memory compression benchmark

BPCells bitpacking compression was compared with a variety of general-purpose compression algorithms. Compression for each data field was measured separately, with algorithms receiving uncompressed binary 32-bit integer inputs. Each algorithm ran compression/decompression repeatedly until at least 2 seconds had elapsed, recording the minimum observed running time. Overall performance was calculated by summing time and file size across the fields present in a file (index/value for matrices, cell/start/end for fragments). Gzip, LZ4 and Zstd were measured using lzbench [31], and the byte-shuffle Blosc variants were measured using the python blosc2 package. Benchmark data at a per-field granularity including additional Blosc filter variants are available in supplemental data. Code available in supplemental code compression/in-memory-compression.

### Bitpacking compression statistics

Bitpacking-compressed files were analyzed for total bytes used per data field and the distribution of bits-per-value in each data field. For the summary statistics shown in Fig S2, the size of metadata (e.g. cell barcodes, gene names) and indexing data are omitted as they are typically negligible in size (*<*2% of total bytes) and/or don’t increase in size as the number of cells is increased. Full bits-per-value distributions and sizes including metadata are available in supplemental data. Statistics for additional matrix non-major axis orderings (sort by number of nonzero entries or random shuffle) are also available in supplemental data. Code available in supplemental code compression/bitwidth-stats.

### RNA matrix storage order transpose benchmark

Input counts matrices for each RNA benchmark dataset were transposed from cell-major order to feature-major storage order. BPCells transpose was performed in a disk-backed manner on compressed BPCells matrix files, while other tools performed the transpose on in-memory data. Memory was limited to 256GB for all tools. Code available in supplemental code rna-timing/matrix-transpose.

### ATAC fragments merge benchmark

Filtered fragment files per sample were prepared in BPCells format or uncompressed 10x fragments.tsv format for each multi-sample dataset. For BPCells, these fragment files were merged and saved to disk in fully genome-sorted order. For GNU sort, fragment files were merged using the parameters LC ALL=C sort -k1,1V -k2,2n -t`$’`*\*t`’`--parallel=4 --merge -S “4G”, then piped to /dev/null to eliminate the overhead of saving uncompressed files to disk.

### CELLxGENE census data preparation

CELLxGENE census long-term supported version 2024-07-01 was downloaded in TileDB-SOMA format from CZI’s hosted S3 files. As BPCells does not directly read the TileDB-SOMA format, the raw counts matrix was converted into BPCells format in 100k cell chunks, along with other metadata necessary for downstream benchmarking. Code available in supplemental code cellxgene/00 convert bpcells chunks.

### CELLxGENE census storage benchmarks

The census data matrices were subset to only include unique cells (marked as is primary data in the census). TileDB configuration followed the settings used in the cellxgene census builder package [1] with buffer sizes increased from 1GB to 4GB on the server, and to 2GB on the laptop. Although cellxgene census builder uses a relatively high Zstd compression level of 9 for integer data and 13 for normalized data, we found that lowering compression levels to 1 saved only 4-12% of write time at a cost of 10-18% larger files, so we use cellxgene census builder default settings (see supplemental data for compression level 1 data).

Normalized data matrices were calculated while sub-setting from the raw counts matrix, following the existing census defaults. For the 0.3% of cells collected with assays where read counts scale with gene length, counts were first divided by gene length (feature length metadata column). Then, counts for each cell were scaled to sum to 1.

For subsetting benchmarks, 10 index selections were randomly generated for each condition (axis, subset size, random/sequential). Each tool was limited to 10 minutes per condition, after which the running selection was allowed to finish but any remaining selections would be skipped to prevent excessive time usage on larger subset sizes. All benchmarks were run with 16 threads on the laptop and 32 threads on the server. Additional performance data subsetting normalized data matrix variants is available in supplemental data. Code available in supplemental code cellxgene/01 subset unique cells and cellxgene/02 matrix slicing.

### CELLxGENE census gene statistics and PCA benchmarks

Mean and variance statistics were calculated per-gene on the subset raw counts matrices using BPCells::matrix stats and cellxgene census.experimental.pp.mean variance. TileDB configuration was taken from the cellxgene census builder package, maintaining default 1GB buffer sizes and 16/32 threads on laptop/server respectively. Additional results with timing for normalized matrices and calculating per-cell statistics are available in supplemental data. Code available in supplemental code cellxgene/03 mean variance.

PCA results were calculated from the feature-major unique-cell matrix with normalizations as described above in “CELLxGENE census storage benchmarks.” On the laptop, 32GB of swap space was enabled to allow for transient memory usage spikes. Code available in supplemental code cellxgene/04 pca.

## Data and code availability

BPCells code, documentation, and open source development is hosted at https://github.com/bnprks/BPCells. Supplemental code for this paper along with supplemental data is hosted at https://github.com/GreenleafLab/BPCells paper. The container image used for benchmarking is hosted at Zenodo along with archival snapshots of the code repositories: https://doi.org/10.5281/zenodo.15066174.

## Acknowledgments

We thank members of the Greenleaf lab for discussion and advice, particularly S. Jessa, B. Liu, and I. Abdi. We also thank Herv’e Pag’es for providing feedback on our DelayedArray benchmarking methodology. Thank you to everyone who contributed to BPCells features that were not bench-marked in this work, with special thanks to I. Abdi. Detailed code contributions are listed at https://github.com/bnprks/BPCells/graphs/contributors. This work was supported by NIH grant R01HG013317. B.P. received support from the JIMB training program funded by NIST during the early development of this work. W.J.G. is an Arc Institute Innovation Investigator.

## Supplemental Discussion

### 1 BPCells performance model

While BPCells utilizes disk-backed streaming algorithms to drastically reduce memory usage compared to in-memory tools, the memory usage of certain operations still scales linearly with the number of cells in a dataset (Fig 1b). BPCells’s memory-usage model assumes there is sufficient memory available to store at least two copies of the principal components matrix. Assuming 50 principal components stored as 32-bit floats, this is equivalent to 100 numbers per cell (0.4GB of RAM per million cells). Downstream tools (e.g. clustering) will already require enough memory to store a single copy of the principal components matrix in memory, and BPCells assumes at least twice this amount is available in order to permit transient copies of the principal components matrix (e.g. in order to transpose the principal components matrix, or to convert from a C++ matrix to an R or Python matrix object). Therefore, BPCells focuses on disk-backed streaming algorithms that can be performed below this memory limit.

From a performance perspective, the simplest form of matrix transformation operation in BPCells is a single-argument mathematical function like log1p. For any gene *g* and cell *c*, the matrix entry *x_gc_* can be transformed as 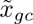 = log1p(*x_gc_*) = log(*x_gc_* + 1). The log1p function has two useful properties for a streaming compute operation:

- *Order independence*: data can be transformed regardless of the order that data points are streamed.
- *Sparsity preserving*: if *x_gc_* = 0, then the transformed value 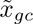 is also 0.

#### Order independence

Order independence means that there are no requirements for the underlying storage ordering for files on disk, so the operation can work efficiently on any input file. The only requirement for order independence is that the transformation function can be expressed as 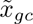 = *f* (*x_gc_, θ_g_, θ_c_*) where *θ_g_* and *θ_c_* are a small set of per-gene or per-cell parameters such as per-gene variance or per-cell read counts to scale the input data. As the number of parameters in *θ_c_* is typically well below 50, the memory usage to store transformation parameters is typically well within the assumed memory budget for BPCells even when chaining multiple transformation operations. An example of a non-order-independent transformation is the rank transformation used for the Wilcoxon rank sum test, since the sorted rank of a matrix entry *x_gc_*_1_ depends on the values of *x_gc_*_2_ and all the other cell measurements for the gene. These transformations can also typically be performed within the BPCells budget, as matrix entries can be buffered to be processed a single cell (column) at a time, which is at worst about as expensive as a transformation that requires 1 parameter per cell. Because the rank transform is not order independent, the Wilcoxon rank sum test sometimes requires first creating a re-ordered copy of the input matrix file, though this re-ordered file can be re-used across multiple order dependent operations.

#### Sparsity preservation

Sparsity preservation can be important for the speed of downstream computations, since losing sparsity can dramatically increase the number of matrix values that must be operated on. All matrix-consuming operations listed in Table S4 are implemented in a sparsity-aware fashion in BPCells, such that the computational cost scales with the number of non-zero matrix entries regardless of how many entries might be zero. In a typical scRNA-seq matrix, only about ^-^7% of matrix entries are non-zero, and an even smaller non-zero percentage is typical in scATAC-seq peak and tile matrices (Fig S2e). Applying additional transformations and calculations after a non-sparsity-preserving operation (even one as simple as 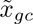 = *x_gc_* + 1) can result in ^-^15x worse performance compared to a sparsity-preserving operation with otherwise similar computational cost.

#### Sparse matrix-vector multiply

Even though non-sparsity-preserving operations generally incur a performance penalty, in some cases there are mathematical rearrangements available that can reduce this cost. This is especially relevant for matrix-vector multiply operations which form the basis for PCA, as mean-centering data is a common normalization prior to PCA that is not sparsity preserving. With some mathematical simplifications, it is possible to do a sparse matrix-vector multiply with a simple pre- or post-processing step which is much faster to compute. For example, if we are adding a per-gene constant to a sparse input matrix *X*, our normalization operation would be 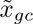 = *x_gc_* + *a_g_*, where *a* is a column vector of per-gene constants. In matrix notation, we can define a matrix *A* = *a*1*^t^* where 1*^t^* is a row vector of all ones, then express our normalized matrix as *X* + *A*. Then to right-multiply by a vector *v*, we have (*X* + *A*)*v* = *Xv* + *Av* = *Xv* + *a*1*^t^v* = *Xv* + *a* ∑*_c_v_c_*. *Xv* is a sparse matrix multiply which is efficient to calculate, and *a* ∑*_c_v_c_* is fast to calculate from the input vectors. A similar simplification approach can work for left multiplies and some other linear but non-sparsity-preserving transformations of a matrix (Table S3).

#### scATAC-seq fragment data

Single cell ATAC-seq matrices share the same performance considerations as scRNA-seq matrices, but handling scATAC-seq fragments is generally simpler. The BPCells fragment data model assumes genome-sorted ordering, so there is no problem of order-dependent operations and fragment data does not have an equivalent of non-sparsity-preserving operations. Therefore, BPCells fragment operations share the same efficiency characteristics of order independent, sparsity preserving matrix operations.

### 2 BPCells programming model

The BPCells implementation relies on a few key C++ and R interfaces in order to allow disk-backed streaming operations to be easily and efficiently combined in a modular way. In C++, the MatrixLoader and FragmentLoader classes define a standard set of interfaces for matrix and fragment objects respectively, which allow disk-backed operations to be freely recombined with each other regardless of the underlying file format or set of operations being performed.

#### C++ matrix interface

MatrixLoader objects (defined in MatrixIterator.h) represent a sparse column-major matrix whose values can be read incrementally. At the C++ level, all matrices are treated as column-major, so a row-major feature×cell matrix file is processed in C++ as if it were a column-major cell×feature matrix with no loss of efficiency. The loading functions load chunks of non-zero entries from within one column at a time, and after loading a data chunk, the MatrixLoader can either load the next non-zero entries from the same column, or skip to reading another column. Frequently, the non-zero entries within a column are ordered by row index, but this is not a strict requirement. When loading a chunk, the MatrixLoader object determines the number of entries to load and provides the data to the reader as raw pointers to row and value arrays. These arrays can be freely modified by the reader until the next chunk of data is loaded.

In addition to the data loading functions, the MatrixLoader interface includes basic metadata functions for the number of rows/columns and names of rows/columns, as well as core math functions for left and right multiply with a dense vector or matrix, and per-row or per-column statistics calculations (sum, mean, variance). These math functions generally use a standard fallback implementation that utilizes the data loading functions to calculate the desired statistics, but they can be specialized to improve performance, such as for faster matrix-vector multiply on mean-centered matrices as described in “Sparse matrix-vector multiply.”

#### C++ fragment set interface

The FragmentLoader interface (defined in FragmentIterator.h) is broadly similar to the MatrixLoader interface. Fragment chunks are loaded in order by start coordinate from one chromosome at a time, and FragmentLoader objects that support seek operations can also skip to an arbitrary chromosome and base pair coordinate at any time (i.e. when using a BPCells file format as input). Performing a seek followed by repeated loads will read all fragments with a start or end coordinate after the seek position, though early loads might also include some fragments with start and end coordinates that fall before the seek position. In addition to loading functions, the FragmentLoader interface contains metadata functions to retrieve chromosome and cell names/counts.

#### Combining objects in C++

For an example of how these loader interfaces are combined in practice, consider the case of loading a raw counts matrix from disk, subsetting the rows, then dividing each column by a constant (e.g. total reads per cell). This will create StoredMatrix, MatrixRowSelect, and Scale objects respectively, all implementing the MatrixLoader interface. Each object contains a pointer to its input matrix loader object, along with C++ objects for file handles, index selections, and scaling constants respectively. Data loading is performed in a “pull” configuration, so a reader would first request data from the Scale object, which would in turn request data from the MatrixRowSelect which would request data from the StoredMatrix object. The StoredMatrix would load data from disk into memory, passing row and value pointers back to MatrixRowSelect along with the number of loaded non-zero entries. Then MatrixRowSelect would modify the row and value arrays to filter out non-zero entries that are not within the chosen row subset and to update the row index to match the index of the output array. Finally, MatrixRowSelect would pass those same row and value array pointers with the new (shortened) chunk size to Scale, which would divide the numbers in the value array by the correct constant for the current column being loaded. It is not required that each object re-use the row and value arrays given as input, though this is frequently done for efficiency purposes. Functions that consume a streaming matrix or fragments object without producing a matrix/fragments object as output are the final stage in all such operation pipelines, and are not themselves part of the MatrixLoader or FragmentLoader C++ class hierarchies.

#### R interface

The R interface provides IterableMatrix and IterableFragments objects and subclasses that mostly have a one-to-one correspondence with C++ subclasses of MatrixLoader or FragmentLoader, with a different subclass for each input file type or transformation operator that BPCells supports. IterableMatrix and IterableFragments objects do not directly store pointers to C++ objects, but rather store the pipeline structure and metadata which can be used to construct the functional C++ loader objects on demand (e.g. file paths and per-row/per-column parameters). When a standard R function such as log1p(), multiplication/addition (*/+), or subsetting ([]) is performed on an IterableMatrix or IterableFragments object no immediate data loading is performed, but the R object is updated with a new outer layer to specify the next operation to perform. Then, once a matrix or fragment consuming function is called (e.g. rowMeans(), svds()), the corresponding C++ objects are constructed and passed to the consumer C++ function which then returns a concrete result to R. This lazy execution strategy has the benefit of minimizing the time that resources such as open file handles are used, as there are no long-lived C++ objects.

### 3 Low-level performance considerations

#### Optimizing for CPU cache usage

Getting data chunks to fit in CPU caches without having to read only from main memory is desirable for performance, but hard to implement for disk-backed libraries like DelayedArray and Dask that are implemented in scripting languages. The performance of CPU cache is much faster than RAM: on contemporary CPUs, a load from the fastest L1 cache typically has over 50x less latency compared to a load from RAM, and L1 cache has bandwidth of 100s of GBs per second *per-core*, compared to 10s of GBs per second of RAM bandwidth that must be split across all cores. BPCells sizes its data load chunks to generally be under 10KB large, which fits well within L1 data cache size (32-48KB per core for our benchmark laptop, and 32KB per core for our benchmark server). This is possible because (1) BPCells uses C++ to minimize per-chunk processing overhead, and (2) BPCells sizes chunks based on the number of non-zero matrix entries to maintain predictable memory footprints.

Dask and DelayedArray are not able to use such small chunk sizes due to higher overheads from performing their data flow coordination in R or Python respectively, rather than C++ as BPCells does with its MatrixLoader and FragmentLoader objects. For example, the “Dask Best Practices” documentation estimates a per-task overhead between 200*µ*s and 1ms, the Scanpy Dask representation recommends a block size of 100k cells, and DelayedArray defaults to a 100 MB block size. These block sizes are 4+ orders of magnitude larger than the comparable values for BPCells. The primary downside of these large chunk sizes is that data does not fit in cache, resulting in worse data read/write performance which is particularly problematic given the low arithmetic intensity of most transformation operations. Secondarily, there are potential problems of system memory bandwidth limitations in multi-core workloads with longer pipelines of delayed operations, as larger-than-cache chunks mean that system memory bandwidth is consumed for every data transformation operator within a pipeline. The BPCells small chunk sizes mean that most data transformations are operating on in-cache arrays, meaning that intermediate operators in a pipeline can consume little-to-no RAM bandwidth.

#### SIMD-accelerated operations

BPCells makes extensive use of SIMD-accelerated operations throughout its implementation. SIMD instructions operate on multiple values at a time, allowing 4, 8, or even 16 values to be calculated with a single CPU instruction. BPCells uses the Highway library [32] to provide cross-architecture SIMD implementations with runtime feature detection to select the most advanced SIMD instruction generation available on the user’s CPU. SIMD instructions are core to the design of BP-128 bitpacking, and they can also provide substantial speedup for mathematical transformations like log1p or SCTransform Pearson residuals [33] (typically about 4-8x performance improvement for AVX2 instructions compared to scalar). To facilitate SIMD acceleration, the MatrixLoader and FragmentLoader interfaces return each data field in a separate contiguous array, sometimes referred to as a columnar memory layout.

Not every operator benefits substantially from SIMD-acceleration, but it can be a very impactful tool to speed up key performance bottlenecks.

### 4 High-level performance considerations

In addition to low-level performance optimizations which target hardware-level efficiencies, BPCells employs high-level optimizations that take place on pipelines of delayed operations in order to minimize compute costs.

#### Subset reordering

A frequent pattern in single-cell matrix code is to perform several normalization steps followed by subsetting to a smaller set of variable genes, or even individual genes for data visualization. In the default programming model, operations are performed in the same order that they are called by a user, which means that BPCells would end up normalizing matrix entries at the start of the pipeline only to discard many of them at a later step. Instead, BPCells re-orders the operation pipeline to put subsetting as early as possible in order to reduce wasted computation. In database terminology, this is an example of predicate pushdown. Nearly all matrix operations in BPCells are compatible with subset reordering, to the point that subsetting a normalized peak matrix can often be pushed all the way to filtering raw fragments loaded from disk before overlap calculations are even performed. All order-independent matrix operations can support subset reordering, and even some order-dependent operations support subset reordering. Rank transforms are the main exception as subsetting prior to ranking will result in different outputs.

#### Parallelization

Multi-threading parallelization in BPCells is performed at a high level to simplify the overall implementation and minimize required communication between threads. The matrix concatenation operators primarily coordinate multi-threading by implementing parallelized versions of MatrixLoader math functions such as matrix-vector multiply. When a user wants to execute a parallel operation on a single matrix, BPCells splits the matrix along its major axis into blocks via subsetting operations (by default 4 times the number of threads), then re-concatenates the blocks with a concatenation operator set to use the desired number of threads. In combination with the subset reordering optimizations and ability to efficiently subset disk-backed matrices along the major axis, this results in each thread loading a subset of the input matrix file, then performing all data decompression, normalization, and aggregation operations itself before the final results from each matrix block are eventually combined into the final output by the concatenation operator. One notable limitation of the existing parallelization implementation is that BPCells does not yet support concatenation reordering, so when working with multiple concatenated matrices, users must manually ensure that the concatenation operation is performed as late as possible in the pipeline. This limitation is likely to be lifted in future software versions.

### 5 Comparison with other software packages

These comparisons reflect the author’s best understanding of the properties of other libraries as of the software version cutoff date for this paper (see Methods), with the aim of helping to contextualize BPCells relative to well-known tools. Although the comparisons focus on design choices that are likely to remain stable over time, details are subject to change as new software versions are released.

#### DelayedArray and Dask

- **Commonalities**: BPCells, DelayedArray, and Dask all form a pipeline of logical operations, perform some high-level optimizations, then execute disk-backed algorithms by operating on matrices one chunk at a time. In general, memory usage should be low for all tools but speed and range of supported operations are differentiating factors.
- **Sparse vs Dense matrix support**: Dask and DelayedArray were originally built to operate on dense matrices and certain operations might require falling back to (slower) dense implementations when a newer sparse implementation is not available. All BPCells streaming operations utilize a sparse matrix representation.
- **Implementation language**: Dask and DelayedArray use Python and R respectively to implement the communications layer between operators. This means new operators can utilize any code in the R or Python library ecosystem, but results in higher overhead compared to the BPCells C++ implementation.
- **Operator pipeline representation**: BPCells and DelayedArray model data pipelines as trees of operators, where any operation might have more than one input but its output can only be used once. Dask supports arbitrary directed graphs of operations and can compute multiple outputs at a time.
- **Storage order awareness**: All operations in BPCells are storage-order-aware, as the BPCells C++-level interfaces treat all matrices as column-major and all fragments as genome-sorted. This allows all operators to optimize based on a shared data ordering and consequent set of efficient seek/subset operations. The core Dask and DelayedArray interfaces take a relatively input-agnostic approach, which makes it easier to support input matrix formats that utilize blocks or tiles (e.g. TileDB), but can make it more complicated to optimize operations which may have to deal with a wide range of input data orderings.
- **Modifying storage order of disk-backed data**: BPCells provides fast disk-backed matrix storage order transposition, which enables users to switch between sparse matrix storage orders to achieve optimal performance for a given operation (see, Figs S1d, S3b). Dask integration in Scanpy and AnnData does not appear to support writing sparse matrices with a different storage order from the input. In DelayedArray, HDF5Array::writeTENxMatrix() supports disk-backed storage order transposition, but with a slow algorithm with apparently quadratic time complexity.
- **BPCells-unique features**: BPCells introduces high-performance matrix storage formats, and also supports scATAC-seq fragment operations which are not addressed by Dask or DelayedArray.

#### TileDB

- **Tile structure**: TileDB supports blocks (or tiles) which can be configured to an arbitrary number of rows and columns. The BPCells sparse matrix format could be thought of as similar to either a single TileDB sparse tile, or a TileDB matrix where tiles are (1 × *n*) or (*n* × 1). BPCells can mimic a similar block structure as TileDB via matrix concatenation operators like with the feature-major 44M cell PCA matrix demonstration, but this involves one matrix file for each block.
- **Mutability**: TileDB supports updating the values in existing matrix files along with version history. BPCells matrix files can only be written once, as single cell data matrices tend not to change once a physical experiment is completed. Appending new data can be simulated in BPCells by writing a new matrix which is concatenated at runtime.
- **Data model**: BPCells matrix files support only 2D sparse matrices, and fragment files support only chromosome, start, end, and cell data for each fragment. The only metadata support is a single name for matrix rows/columns or chromosome/cell names in the case of fragments. TileDB allows multi-dimensional arrays, sparse or dense arrays, and each matrix entry (cell in TileDB terminology) can store multiple types of values. TileDB can also be used for storing tabular data such as for per-cell metadata in the TileDB-SOMA model.
- **Integrated computation**: BPCells provides efficient disk-backed computations along with its file format. TileDB provides just a data format.
- **Multi-threading**: TileDB supports multi-threaded reads and writes. Every BPCells thread interleaves both data loading and computation, so multi-threaded matrix computations result in multi-threaded reads. BPCells writes are single-threaded per file, though as we demonstrate in Figs 3b+S4a, the BPCells on-the-fly concatenation operations make it possible to split a large matrix into multiple files that can be quickly written in parallel.
- **Compression options**: TileDB allows a wide range of configurable filters and compressors including gzip, Zstd, and LZ4 which are similar to those benchmarked in Fig 2e+j and Fig S3a. TileDB does support a bitpacking-style filter (TILEDB BIT WIDTH REDUCTION), but it seems to allow only 8, 16, 32, or 64 bits and not to utilize SIMD acceleration. BPCells uses only BP-128 based bitpacking compression with a fixed set of per-field filters which are integrated into the SIMD-accelerated BP-128 code (i.e. operator fusion). Additionally, BP-128 supports any integer bitwidth from 1-32 which helps to provide better space savings for data that requires a non power-of-two number of bits.
- **Configuration**: BPCells provides minimal configuration options, mainly just to enable/disable compression and store data in row-major/column-major order. In general, BPCells aims to provide one or two simple options that are high performance across most single cell analysis use-cases. TileDB supports many domains beyond single-cell genomics, so it provides a large number of per-file configuration options as well as runtime configuration options which may have to be tuned for optimal performance in any given use-case.

#### ArchR and SnapATAC2

- **Commonalities**: BPCells, ArchR, and SnapATAC2 all support fragment-level scATAC-seq operations, disk-backed peak/tile matrix creation, and ATAC-seq quality control metric calculations such as TSS enrichment.
- **Fragment storage layout**: BPCells stores fragments in genome-sorted order. ArchR and SnapATAC2 group fragments by chromosome and cell, then store start and end coordinates for each fragment. We highlight the performance advantages of the BPCells approach in Fig 1f+g. This decision also means that BPCells produces peak/tile matrices in feature-major order, while ArchR and SnapATAC2 produce them in cell-major order.
- **Modular streaming operations**: BPCells provides disk-backed streaming operations that can be freely recombined to create more complex normalizations and computation pipelines that can run in a single pass over the data. In ArchR and SnapATAC2, any disk-backed operation must execute completely and save its result back to disk before another operation can use its output. For example, BPCells provides a wide range of fragment filtering and manipulation options (Table S5) which can be applied on-the-fly without having to modify or re-write data on disk, whereas ArchR and SnapATAC2 require specifying parameters like Tn5 offset shifts at data import time with no option to adjust without re-importing raw data.
- **Full-precision disk-backed PCA**: BPCells can perform fully disk-backed PCA and other SVD-based dimensionality reductions. ArchR and SnapATAC2 require loading the feature×cell matrix into memory to perform full-precision dimensionality reduction. ArchR supports using a subset of cells to reduce memory usage for dimensionality reduction, and SnapATAC2 supports a randomized approximate disk-backed dimensionality reduction.

#### Scanpy and Seurat

- **Commonalities**: Scanpy, Seurat, and BPCells all enable single cell matrix normalization, PCA, and marker feature testing.
- **Disk-backed support**: All BPCells operations are disk-backed. Seurat and Scanpy have optional integrations with disk-backed computation libraries (BPCells as a back-end for Seurat, and Dask as a back-end for Scanpy). Not all features in Seurat and Scanpy support disk-backed execution, and all of Scanpy’s disk-backed operations with Dask appear to be in experimental status currently.

### 6 PCA performance considerations

Because PCA is one of the only operations in BPCells that requires multiple matrix passes to complete, it offers a new set of performance considerations around whether to make temporary intermediate matrix copies on disk. For the sake of this performance analysis, we assume a PCA procedure where a raw counts matrix is (1) optionally subset to variable features, (2) log-normalized, (3) z-score normalized, then (4) the principal components solver performs repeated matrix-vector multiplies with the normalized matrix in order to compute the top principal components. Recall that the z-score normalization step can be combined with the matrix-vector multiply operation at little cost, as outlined in “Sparse matrix-vector multiply,” so we consider it a part of the matrix-vector multiply operation for the remainder of this section.

Because PCA typically uses over 100 matrix-vector multiplies, it can sometimes be beneficial to make an intermediate matrix copy to avoid repeating filtering and normalization calculations during each matrix-vector multiply. We consider three main strategies:

- **No intermediate**: Perform filtering and log-normalization on the fly during each matrix-vector multiply.
- **Integer intermediate**: Save a counts matrix subset as an intermediate copy, eliminating the need to load and filter out non-variable features during each matrix-vector multiply. Log-normalization is still repeated for each multiply.
- **Normalized intermediate**: Save a log-normalized, variable features matrix subset as an intermediate copy. This results in only having to compute the log-normalization once, reducing the cost for all future matrix-vector multiplies during PCA.

For the comparative benchmarks in Fig 1b-d, we used the “normalized intermediate” strategy, as subsetting from 20k-60k genes down to 2k variable genes means that the size of the intermediate normalized matrix would not be too large. On our benchmark datasets, the 2k variable gene subset on average had just 8% of the nonzero entries as the full matrix, so intermediate storage costs were relatively low despite the inability to apply bitpacking compression on the values of the normalized matrix. We included benchmarks of the other strategies in supplemental data.

For the 44M cell atlas-scale demonstration in Fig 3f+h, we compared the “no intermediate” strategy with the “normalized intermediate” strategy, notated as “compressed” vs. “uncompressed” inputs. In this highly multi-threaded benchmark, disk bandwidth was a scarcer resource than compute, so the smaller file sizes of the compressed inputs outweighed the reduced computational requirements for the normalized but uncompressed inputs. For the randomized PCA algorithm, only 6 matrix passes were required so it was fully compute-bound and there was little-to-no difference in performance based on compressed or uncompressed inputs.

#### Operating system file caching

One confounding factor present in most of our disk-backed benchmarks is the presence of operating system file caching, where the contents of frequently-used files may be loaded into RAM in order to allow much faster read speeds. In the case of PCA benchmarks, the input file will be read multiple times and the Linux operating system can cache the entire input file in RAM (assuming sufficient memory capacity is available). Since this memory usage is done at the level of the operating system, it is not attributed to our benchmarked processes but nevertheless can help speed up reads from disk. Thus, certain disk-backed operations will run faster on systems with more RAM even if they can still successfully run on a system with less RAM.

We can observe the effects of file caching most clearly in the server PCA results for our 44M cell benchmark in Fig 3h. Using a compressed input compared to an uncompressed input resulted in a whopping 9x faster execution speed, compared to just a 1.5x change on a laptop. This is because the server’s RAM could fully hold the 117GB compressed input matrix, allowing average read speeds of 6.3GB/s due to file caching. By contrast, the 765GB uncompressed matrix was too large to cache in RAM, and averaged a lower 4.6GB/s read speed from disk. On the laptop where neither matrix fit in memory, these numbers were reversed as both matrices were read from disk and the laptop with compressed input was also more heavily compute bottlenecked (0.9GB/s for compressed and 3.9GB/s for uncompressed inputs respectively).

The effects of file caching are also apparent in the comparative normalization and PCA benchmarks, though likely not in a way that seriously affects the comparisons in Fig 1b-d. With the plotted “normalized intermediate” strategy, we observed average read speeds uniformly below 2GB/s for single-threaded execution and below 4GB/s for 2-threaded execution, which are well within the bandwidth capabilities of the server SSDs whether or not OS file caching was in effect (see Methods for hardware details). In the multi-threaded benchmarks shown in Fig 1d we saw clear effects of file caching, with average file read speeds in some datasets going beyond 20GB/s when 16 threads were used for PCA – well above the physical capabilities of the server SSDs. In most real-world use cases, it’s likely that the user’s computer would have sufficient RAM to cache these files. The largest of the intermediate matrix files was just 7.5GB in size, and in most cases the intermediate matrix files were smaller than the already observed maximum memory usage for the PCA processes. Therefore, when subsetting to 2k variable features prior to PCA, we expect that users will also have sufficient spare RAM to cache the intermediate matrix files and achieve speedups for multi-threaded execution without bottlenecking on disk bandwidth like we observed for the 60k gene, 44M cell server benchmark discussed above.

One minor difference in the execution of the “normalized intermediate” strategy between the 44M cell benchmark and the comparative benchmarks in Fig 1 is that in the 44M cell benchmark no compression at all was applied to the “uncompressed” variant, while in Fig 1 the staged file still had the row index data compressed as usual, just the normalized value data was left uncompressed. Compressing just the row index data still resulted in an overall geometric mean 1.6x compression ratio across the tested datasets, which helps to moderately reduce disk bandwidth when the files are not cached and also makes it more likely for the intermediate matrix data to fit in the operating system file cache.

### 7 Peak and tile matrix creation algorithm

The BPCells peak and tile overlap matrix algorithms are markedly faster than the alternatives from ArchR and SnapATAC2 as illustrated in Fig 1e-g. Across all the peak and tile matrices calculated for benchmarking, BPCells averaged 33M overlaps tabulated per second for peak matrices, and 9M overlaps per second for tile matrices (single-threaded). To accomplish this, BPCells uses a linear scan algorithm that leverages the known sorted order of input fragments, focusing optimization efforts on efficient CPU utilization via cache and branch predictor friendly design choices.

In this section, we describe the algorithms BPCells uses to convert a stream of sorted scATAC-seq fragments into a streaming sparse matrix of dimensions cells×regions, where each matrix entry holds the count of overlaps between a given (cell, region) pair summed across all fragments. We focus first on the peak matrix algorithm, then describe the relevant adaptations required for tile matrix calculations at the end.

The code implementing these algorithms can be found in files PeakMatrix.h, PeakMatrix.cpp, TileMatrix.h, TileMatrix.cpp, and MatrixAccumulators.h.

#### Task specification

- **Fragment inputs**: Fragments are input as a FragmentLoader object, which loads a stream of fragments with (chromosome, start, end, cell ID) in sorted order by (chromosome, start). The input loader can optionally support seek operations which allow skipping to a given (chromosome, coordinate) position such that the following reads will capture all fragments where the end position falls after the given coordinate.
- **Peak region inputs**: A set of peak regions given as (chromosome, start, end). Peaks can be any size and are allowed to be arbitrarily overlapping or nested. Peaks are assigned numeric IDs based on their index when the list of peaks is sorted by (chromosome, end).
- **Output matrix**: The output matrix is a MatrixLoader object which gives a stream of (row, column, value) non-zero matrix entries, where the row is a numeric cell ID, the column is the peak index ordered by (chromosome, end), and the value is the total count of overlaps. Because fragments are read in genome-sorted order, overlap calculations are completed one peak at a time so the output matrix is dimensions cells×features (i.e. peaks) in feature-major order.
- **Overlap calculation options**: BPCells offers three options for how fragment-peak overlaps are calculated:

– *insertions*: Start and end coordinates for each fragment are separately overlapped with each peak. If both the start and end for a fragment fall within a peak it counts as two overlaps. If the fragment fully spans the peak such that the fragment start is before the peak start and the fragment end is after the peak end then no overlap is counted. This option is the default for scATAC-seq fragments where the fragment ends represent the sites of genomic accessibility.

– *fragments*: Similar to the “insertions” option, but if both the start and end for a fragment fall within a peak it only counts as one overlap.

– *overlaps*: Similar to the “fragments” option, but an overlap is also counted if the fragment fully spans the peak such that the fragment start is before the peak start and the fragment end is after the peak end.

#### Overlap detection

The first task in peak matrix creation is detecting overlaps between fragments and peak regions. The BPCells formulation of overlap detection takes in a stream of sorted fragments (chromosome, start, end, cell ID) and outputs overlap data points (cell ID, peak ID, overlap count). Depending on the overlap calculation option used the overlap count can be 1 or 2. Each overlap data point corresponds to exactly one input fragment, but the reverse is not true since one fragment can overlap more than one peak.

To minimize the required number of overlap calculations, BPCells conceptually divides peaks into three states which they pass through sequentially:

1. **Inactive** peaks have a start coordinate higher than any observed fragment end coordinate so far, and thus can’t produce overlaps yet.
2. **Active** peaks are the only ones directly checked for fragment overlaps, as the current chunk of fragment data may contain start or end coordinates within the peak. The active peak set is explicitly stored in a C++ vector.
3. **Completed** peaks have an end coordinate lower than all the remaining fragment start coordinates, so they cannot produce any more overlaps and the corresponding peak matrix columns can be output.

Peaks will transition from inactive to active in order by start coordinate, and will transition from active to completed in order by end coordinate. Therefore, BPCells stores the ordering of peaks sorted by start and end coordinates so it can easily track the next peak that could potentially transition from inactive to active, or from active to completed. For the inner loop of overlap detection, BPCells loads a chunk of fragments (typically around 1,000), then subdivides these into blocks of 128 for overlap processing. For each block of 128 fragments, BPCells performs the following steps:

1. Check the maximum end coordinate of any fragment in the block, then if the maximum end coordinate is larger than the start coordinate of any inactive peaks, transition them to active.
2. For each peak in the active set:

(a) Calculate overlaps with the block of 128 fragments using SIMD-accelerated code.
(b) If the largest start coordinate of the block is larger than the peak end coordinate, remove the peak from the active set.

If peaks have been moved from the active set to the completed set after processing a chunk of fragments, then BPCells will potentially pause loading fragments to perform overlap aggregation and outputting peak matrix entries.

#### Overlap aggregation

Because multiple fragments from a cell might overlap the same peak, there may be multiple (cell ID, peak ID, overlap count) data points that correspond to a single non-zero entry in the output cell×peak matrix. To deal with this, BPCells uses an overlap aggregation data structure which buffers overlap data points then merges overlaps from the same cell and region to produce non-zero entries for the output peak matrix. This overlap aggregation is a key step in high performance peak matrix creation, and several potential solutions such as hash tables and per-region vectors were considered before arriving at the solution described below.

First, as each overlap data point is found during overlap detection, it is appended to a single dynamically grown buffer of (cell ID, peak ID, count) data points. When this buffer fills up or output matrix data is required, a compaction operation takes place which first sorts the buffer by (peak ID, cell ID), then merges adjacent entries with matching (peak, cell) values by summing the counts. If the buffer is too full after compaction, then it is grown multiplicatively. After compaction, any buffer entries corresponding to peaks with “completed” status can be output directly as nonzero entries in the peak matrix, with the cell ID corresponding to the row index, peak ID corresponding to the column index, and count as the matrix value. After values have been read from the output MatrixLoader object for the peak matrix, they can be removed from the buffer.

#### Key performance considerations

The main optimizations in peak matrix calculations focus on cache-friendly memory access patterns and avoiding branch mispredictions:

- In overlap detection, the inner loop of SIMD-accelerated overlap checking minimizes branches based on the outcome of fragment overlap calculations, which helps to avoid branch misprediction delays.
- In overlap detection, if the active peak set ever becomes empty the fragment loader object can perform a seek operation to advance the input stream to fragments that could potentially overlap the next peak ready to activate.
- When an overlap is detected, adding it to the overlap aggregation buffer is a cheap, cache-friendly append to the end of the buffer.
- During compaction for overlap aggregation, sorting is done via a least significant digit radix sort which has cache-friendly memory access patterns and uses a branching pattern that minimizes branch mispredictions. When sorting 32-bit integers, we observe the radix sort runs about 10x faster than C++ std::sort() while being relatively simple to implement.

From an algorithmic complexity perspective, this algorithm provides few guarantees in worst-case pathological examples. If the length of fragments is much longer than the spacing between adjacent peak regions, the set of active peaks may become very large and increase the number of overlap checks required during overlap detection. For example, if the dataset contained one fragment on each chromosome that spanned from the first base pair to the last base pair of the chromosome, the algorithm described above would have quadratic time complexity by having to directly compare every fragment with every peak for overlaps. There are likely adjustments to the described algorithm that could avoid this worst-case time explosion, but we have not implemented them yet in BPCells as such pathological examples are unlikely to ever be seen in practice given the physical constraints of the molecular biology methods used to collect the underlying scATAC-seq data (or even CUT&Tag data).

#### Tile matrix calculation

Tile matrix calculations utilize a very similar approach as peak matrix calculations with a few modifications:

- **Input regions**: Inputs are specified as regions which are subdivided into fixed-width tiles with tile widths specified on a per-region basis. Regions might be full chromosomes tiled at 500bp resolution, or could be smaller regions of interest tiled at higher resolution for visualization purposes. Output tile IDs are assigned sequentially by genomic position and input regions must not overlap so that all tiles from a single region have consecutive tile IDs.
- **Overlap calculation options**: Only the “insertions” and “fragments” modes are available for tile matrix calculations.
- **Overlap detection**: Overlaps are detected at the granularity of regions, then an integer division is used to determine which tile from within a region a fragment overlaps with. Because the tile width is fixed for each region,

BPCells uses a vectorized version of the libdivide algorithm [34] to translate slow integer division instructions into faster addition, multiply, and bitwise shift instructions.

### 8 Wilcoxon rank sum algorithm

The BPCells Wilcoxon rank sum function takes as input a feature×cell matrix along with cell group assignments (e.g. cell types), where each cell is assigned to a group. Here we will discuss it in the context of gene expression, but the calculations apply equally to chromatin accessibility or any other type of feature. For each gene and group, BPCells performs a Wilcoxon rank sum test comparing the distribution of gene expression values for cells within the group compared to all cells outside of the group (one-vs-rest), eventually calculating a p-value for each (gene, cell group) pair. We omit a detailed discussion of the statistical test as it is well described in many other sources. The code implementing these algorithms can be found in files WilcoxonRankSum.h, WilcoxonRankSum.cpp, ColwiseRank.h, and radix sort.h.

From a computational perspective, the steps are:

1. Load all expression values for one gene across all cells.
2. Sort the expression values to calculate a rank for each cell.
3. Sum the ranks for each cell group, calculate an additional statistic based on the number of tied values, then compute p-values.

Compared to Presto [14] discussed in Fig S1d, BPCells is able to gain efficiency in two main places. First, step (1) can be done efficiently in BPCells by using a feature-major input matrix, and if the original dataset is not available BPCells can convert a matrix from cell-major to feature-major efficiently (Fig S3b). By contrast, Presto operating on a sparse matrix input requires cell-major input then switches the storage order to feature-major in memory prior to running calculations. On the 500k cell test dataset, BPCells runs 2.3x faster than Presto when BPCells has a feature-major input. If we add the time for BPCells to switch storage orders, the performance gap shrinks to 1.5x.

The other likely source of BPCells’s improved performance over Presto is its use of sorting algorithm to compute the rank sums. BPCells uses a radix-based sorting algorithm which is adapted to work on floating-point or integer data. Although radix sort is most commonly described for integers, simple bitwise pre- and post-processing can transform floating-point or signed integer data into unsigned integers that share the same relative sorting order. Compared to Presto which uses the C++ std::sort() function, we find a radix sort has roughly 10x faster performance. This faster sorting algorithm likely accounts for the remainder of the performance advantage compared to Presto, as the remaining steps are not very computationally intensive.

Finally, we note that the BPCells rank transformation actually computes per-feature ranks with an offset added to make the rank of 0 within each feature map to 0 (Table S3). This offset is easily reversed during rank summation, but preserves the matrix sparsity structure to greatly reduce the number of matrix entries that must be explicitly processed during rank summation and tie statistic calculations. Presto appears to use a similar approach, but does not appear to handle the edge case where negative values might be present in the input matrix.

### 9 Matrix storage order transpose algorithm

As discussed in “Data storage ordering,” BPCells must provide an efficient, disk-backed algorithm to switch between row-major and column-major matrix layouts. For an in-memory CSC-or CSR-format sparse matrix, the storage order transpose operation requires a two-pass, linear time counting sort algorithm that relies on RAM’s fast random writes to re-partition data. However, in a disk-backed context random access is much slower than sequential read/writes (even for SSDs), so BPCells takes an external merge sort approach which allows fully sequential writes and random reads in large megabyte-scale blocks. The code implementing this algorithm can be found in StoredMatrixTransposeWriter.h, StoredMatrixSorter.h, and radix sort.h.

The external merge sort first loads (row, col, val) matrix entries from the input MatrixLoader object which are sorted in order of (col, row). BPCells allocates 1GiB of memory to sort 500MiB blocks of matrix entries by (row, col) with a least significant digit radix sort, then writes the sorted blocks concatenated in bitpacked files for row, column, and value using BP128-d1z, BP128-d1, and BP128-m1 encodings respectively in order to maximize effective disk speed and minimize intermediate disk space usage (see BP-128 compression variants; no compression used for non-integer value types). Then, BPCells combines sorted blocks into larger blocks by using a heap to track which input block has the next matrix entry to output. Because input data blocks will tend to span several adjacent columns of the input matrix, the same data block will tend to output several consecutive entries into the output. Therefore, BPCells tracks the second-largest value in the heap to avoid a heap pop/push operation when the maximum data block stays unchanged.

With the default settings of 1GiB sorting buffer and 4MiB minimum read size, BPCells can perform 85-way merges, allowing it to process up to 46GB of uncompressed data in 2 passes, or 3.9TB of uncompressed data in 3 passes. Increased sorting buffer sizes can increase the data sizes which can be processed in 2 or 3 passes.

This algorithm is re-used for efficient MatrixMarket (mtx) file import as there is no guaranteed ordering of non-zero matrix values within a MatrixMarket coordinate file.

### 10 Sorted fragment merge algorithm

Because BPCells requires fragment files to be sorted by genomic coordinate and allows on-the-fly merging of fragment files, it must also implement a fast sorted merge algorithm for fragments data. For this algorithm, BPCells takes as input *N* FragmentLoader objects, producing a single FragmentLoader that merges the sorted fragment streams into a single combined stream. The code implementing this algorithm can be found in MergeFragments.h and MergeFragments.cpp. BPCells uses a hybrid of a heap-based merge and radix sorting. By default, input is loaded in chunks and a value chunks per input is chosen to be at least 2 and up to load size*/*(chunk size ∗ *N*) (where load size defaults to 1024 and chunk size defaults to 32). Given this setup, the merge algorithm performs the following steps:

1. **Load data**: Load chunks per input ∗ *N* chunks of chunk size fragments from the *N* input loaders into a data buffer. BPCells uses a heap over the input FragmentLoader objects to always add the chunk with the smallest start coordinate. BPCells also tracks the running maximum of these smallest start coordinates.
2. **Sort data**: After loading all chunks, perform a least significant digit radix sort to order fragments in the data buffer by start coordinate.
3. **Output data**: Fragments with start coordinates up to the running maximum of smallest start coordinates among added chunks can be safely output then removed from the data buffer as they are known to be in the final sorted order.

This algorithm guarantees that there is at most 1 chunk of fragments left over in the buffer per input FragmentLoader (chunk size ∗ *N* fragments). Therefore, we size the buffer to (chunks per input + 1) ∗ chunk size ∗ *N*, guaranteeing that there is always space to load chunks per input ∗ chunk size ∗ *N* fragments into the buffer for each new loading step. Therefore, the average number of times each fragment must be sorted is at worst (1 + chunks per input)*/*chunks per input.

The initial implementation of fragment merging used a simple heap-based merge loading 1 fragment at a time, but this proved to be slower than the hybrid heap + radix sort design, likely due to reduced heap operations and branch mispredictions in the hybrid design.

### 11 Matrix storage ordering recommendations

Matrix storage in BPCells has two main axes of customization to improve performance for a given application. First is choosing between a cell-major or feature-major storage order, which allows efficient subsetting of individual cells or features respectively. Second is the option to split a large matrix into multiple files as we demonstrate for the 44M cell analysis of the CELLxGENE census, which can help with parallelization in certain circumstances with extremely large datasets. We’ll describe our recommendations and rationale for optimal matrix storage for several use-cases in this section.

#### Neural network training

Recent years have seen a growing interest in training neural network models that can leverage large single cell datasets to learn to predict gene expression profiles for cells. From a data access perspective, training these models requires loading mini-batches of cells, typically random samples of ^-^10-100 cells. For this use-case, cell-major storage orders are an absolute requirement to enable fast random access to small numbers of cells. In Fig 3d we observe queries of 1-100 cells taking roughly half a second to complete. It may be preferable to load larger batches then split into mini-batches to achieve higher throughput, as we observe 10k cells loading in 1 second and 100k cells loading in 3 seconds. In cases where even faster performance is required, compressed matrix data could be stored in host RAM (44M cells should fit in *<*170GB of RAM). Splitting matrices into blocks is not necessary for querying, but may help accelerate file management tasks as described below.

#### General single cell analysis

For general single cell analysis tasks such as running normalization, PCA, and marker feature testing, we recommend storing at least a feature-major counts matrix and optionally adding a cell-major matrix copy to allow optimal performance in all scenarios. The advantages of a feature-major layout are:

- **Efficient marker feature and differential feature tests**: the Wilcoxon test we demonstrate in this paper requires a feature-major storage order to run efficiently, and this would also apply to many other differential testing methods such as those involving generalized linear models (e.g. DESeq2).
- **Visualization of genes/features**: For visualization purposes it is often useful to plot the distribution of gene expression for a small number of genes across all cells, such as in UMAP, violin, or dot plots. Feature-major orderings allow fast subsetting by gene/feature to make such visualizations fast to produce.
- **Improved compression**: Whether using bitpacking compression or general-purpose algorithms, a comparison of Figs 2e and S3a shows that data storage size tends to be smaller with a feature-major storage ordering.

The potential downsides of a feature-major layout are:

- **Slower subsetting by cell**: If a simple subset of a very small fraction of the dataset is required, that will execute faster with a cell-major layout. In practice, this downside can often be mitigated by creating temporary matrix copies for cell subsets of interest (e.g. cell types or batches), or by performing operations that can process multiple cell subsets in a single pass such as calculating pseudobulks.
- **Memory usage during parallel PCA**: When calculating *k* principal components with *t* threads, the BPCells multi-threaded dense matrix multiply operation can require memory of *t* × *k* × # cells when performed on a single feature-major matrix file, which is potentially much larger than the assumed memory budget of 2 × *k* × # cells (see BPCells performance model). Splitting the matrix along the cell dimension into *t* or more chunks can reduce this memory usage to *k* × # cells.

Note that splitting by cell does cause multiplication with a *k* × # features matrix to require *t* × *k* × # features memory, but this is not problematic in single cell datasets where there are generally at most 100k features, but can be more than 10M cells.

#### **>**10M-cell atlas analysis

For extremely large atlas datasets, parallelizing file writing operations in addition to file read/compute operations can become very important. For these cases, we recommend splitting a large input matrix into multiple matrix files each containing a subset of cells, allowing multiple threads to write separate files in parallel. We use this approach in Fig 3 by splitting input matrices into 100K cell chunks, allowing fast subsetting from the duplicated 74M cell CELLxGENE census matrix to 44M unique cells (Fig 3b). If only a single matrix copy can be stored, then splitting a large matrix into 100K-1M cell chunks each stored in feature-major layout is often preferable to storing each matrix chunk in cell-major layout for the same reasons as discussed above in “General single cell analysis.”

Splitting matrices into 100K-cell chunks has side benefits for sequential subsets and appending new cells to a matrix. Having 100K-cell files allows some coarse-grained random access, making it possible to perform reasonably efficient sequential subsets by cell even when the underlying files use a feature-major storage order (Fig S4f+g). Grouping cells in a large atlas dataset by sample and/or cell type can turn many common cell subset queries into sequential subsets, meaning that many cell subsets could be performed with reasonable efficiency even from feature-major storage orderings. In addition, when new cells are added to a large atlas, using 100K-cell chunks means at most one existing matrix chunk must be re-written.

In the case that there is sufficient disk space to store two raw counts matrices, storing both a feature-major copy and a cell-major copy split into 100K-cell chunks is advantageous, as cell-major layouts are still better suited for neural network data loader applications as described above.

### 12 BPCells file formats

Here we outline the technical specifications for the storage systems, bitpacking variants, and sparse and matrix file formats that BPCells introduces. These correspond to the BPCells-specific file formats in the list of all supported file formats from Table S7.

#### Storage back-ends

BPCells supports a variety of storage back-ends which can be selected for performance or portability reasons. For benchmarking, we use the “directory of files” backend, but BPCells can be readily adapted to any storage backend that meets simple interface criteria. The core storage abstraction required for BPCells matrix or fragment files is a set of arrays which can be independently accessed by name, where each array can store numeric or string data types. Numeric arrays must support efficient random access. We also require some mechanism to store a version string for compatibility purposes. The current supported storage backbends implemented by BPCells are:

- **Directory of files**: This is the default storage backend due to its simplicity and high performance. Arrays are stored as binary files within a directory. Numeric array files have an 8-byte header followed by data values in little-endian binary format for integers and IEEE-754 for 32-bit and 64-bit floating point numbers. Header values are 8-byte ASCII text as follows: unsigned 32-bit integer UINT32v1, unsigned 64-bit integer UINT64v1, 32-bit float FLOATSv1, 64-bit float DOUBLEv1. Arrays of strings are stored as ASCII text with one array value per line with no header. The version string is stored as a file named “version” containing the version string followed by a newline.
- **HDF5 file**: This storage backend can be useful for embedding BPCells formats as a group within an h5ad or other HDF5 file. Arrays of numbers are stored as HDF5 datasets using the built-in HDF5 encoding format. Arrays of strings are stored as HDF5 variable length string datasets. The version string is stored as a version attribute on the HDF5 group.
- **R object**: This storage backend is primarily useful for testing, or when bitpacking compression of in-memory data is desired to avoid disk bandwidth bottlenecks. Strings are stored as native R character arrays. Unsigned integers and 32-bit floats are stored in native R integer arrays by bitcasting the R signed integers into the required data types. 64-bit floats are stored in native R numeric arrays. 64-bit integers are stored as doubles in R numeric arrays. This reduces the highest representable value from 2^64^ − 1 to 2^53^ − 1 (about 9 quadrillion), which we do not expect to pose practical problems. Named collections of arrays are stored in R lists (when writing) or S4 objects (when reading). The version string is stored as a string vector named “version” of length 1.

#### BP-128 compression variants

BPCells uses BP-128 compression and variants of it extensively to provide compact storage for arrays of 32-bit integers (Fig 2a+f). The original BP-128 paper by Lemire and Boytsov [6] does not specify a concrete storage layout, and BPCells chooses to split each compressed data stream into 3-4 arrays depending on the associated transformations. As BP-128 works on chunks of 128 integers, input lengths are padded to a multiple of 128 by repeating the last integer in the list. When decompressing data, the length of the output data array is calculated from other uncompressed metadata fields present in the matrix or fragment objects. The variants of BPCells and their corresponding data arrays are as follows:

- **BP-128**: Best for lists of small, non-negative integers. Worst-case overhead: 0.25 bits/integer (0.8%).

– data: stream of bitpacked data, represented as 32-bit integers with the interleaved bit layout as shown in Lemire et al. figure 6 [6]. A chunk of 128 32-bit input integers with *B* bits per integer will be stored using 4*B* 32-bit integers holding the bitpacked data. Explanatory C code for compression and decompression can be found in supplemental code utils/bitpacking-reference-implementation.cpp (little-endian byte order is assumed).

– idx: list of 32-bit integers, where the encoded data for integers index 128*i to 128*i + 127 can be found in data from index idx[i] to index idx[i+1]-1. For lists with 2^32^ (4 billion) entries or greater, idx stores the index modulo 2^32^

– idx offsets: list of 64-bit integers, where the values of idx with indices from idx offsets[i] to idx offsets[i+1]-1 should have i*(2^32) added to them.

- **BP-128d1**: Equivalent to the BP-128* algorithm from Lemire et al. [6] where integers are difference encoded prior to bitpacking. Best for lists of ascending integers. Worst-case overhead: 0.5 bits/integer (1.6%).

– data: Encoding as with vanilla BP-128, but prior to encoding we perform difference encoding, so 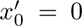, 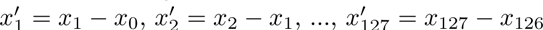

– idx, idx offsets: identical to BP-128

– starts: list of 32-bit integers, where starts[i] is the decoded value for the integer at index 128*i

- **BP-128d1z**: Similar to BP128d1, but with zigzag encoding after difference encoding. Best for lists of close but not fully sorted runs of integers. For fully sorted data, overhead is about 1 bit per integer compared to BP-128d1. See supplemental code utils/bitpacking-reference-implementation.cpp for zigzag encode() and zigzag decode() reference implementations.

– data: Encoding as with BP-128d1, but between difference encoding and bitpacking, the results are zigzag encoded, where *zigzag*(*x*) = 2*x* if *x* ≥ 0, and *zigzag*(*x*) = −2*x* − 1 if *x <* 0.

– idx, idx offsets: identical to BP-128

– starts: identical to starts from BP-128d1

- **BP-128m1**: Similar to BP-128 but with 1 subtracted from all values prior to encoding. Best for lists of small positive integers with many equal to exactly 1.

– data: Prior to bitpacking, 1 is subtracted from all values.

– idx, idx offsets: identical to BP-128.

#### Sparse matrix format

BPCells matrix files are stored in compressed sparse column (CSC) or compressed sparse row (CSR) layouts. For CSC layout, non-zero entries are sorted in ascending order by row within each column and vice-versa for CSR layout. The matrix data and metadata is stored as a named set of arrays using one of the previously described storage back-ends.

Uncompressed matrices consist of the following arrays:

| Name | Type | Length |
| --- | --- | --- |
| **val** | uint32/float32/float64 | # non-zeros |
| **index** | uint32 | # non-zeros |
| **idxptr** | uint64 | # cols + 1 |
| **shape** | uint32 | 2 |
| **row_names** | string | 0 or # rows |
| **col_names** | string | 0 or # cols |
| **storage_order** | string | 1 |

Arrays for uncompressed sparse matrix storage

These arrays have the following contents:

- val: Values of non-zero entries in increasing order of (column, row) position.
- index: index[i] provides the 0-based row index for the value found in val[i] (or column index for row-major storage order).
- idxptr: The indexes in idx and val for the entries in column j can be found from idxptr[j] to idxptr[j+1] - 1, inclusive (or row j for row-major storage order).
- shape: number of rows in the matrix, followed by number of columns
- row names: Names for each row of the matrix (optional)
- col names: Names for each column of the matrix (optional)
- storage order: col for compressed-sparse-column, or row for compressed-sparse-row

Compressed matrices consist of the following modifications:

- val: For unsigned 32-bit integers, we replace val with val data, val idx, and val idx offsets corresponding to a BP-128m1 encoding as described above. The total number of values is already stored as the last value in idxptr. For 32-bit and 64-bit floats val remains unchanged.
- index: We replace the index array with BP-128d1z encoded data in arrays index data, index idx, index idx offsets and index starts

The current version string is of the format: [compression]-[datatype]-matrix-v2, where [compression] can be either packed or unpacked, and [datatype] can be one of uint, float, or double corresponding to 32-bit unsigned integer, 32-bit float, and 64-bit double respectively. In v1 formats, the only difference is that idxptr had type uint32.

#### Genomic fragments format

BPCells fragment files store (chromosome, start, end, cell ID) for each fragment, sorted by (chromosome, start). The coordinate system follows the bed format convention, where the first base of a chromosome is numbered 0 and the end coordinate of each fragment is non-inclusive. This means a 10 base pair long fragment starting at the first base of the genome will have start=0 and end=10. End coordinates are always guaranteed to be at least as large as start coordinates.

Uncompressed fragment data is stored in the following arrays:

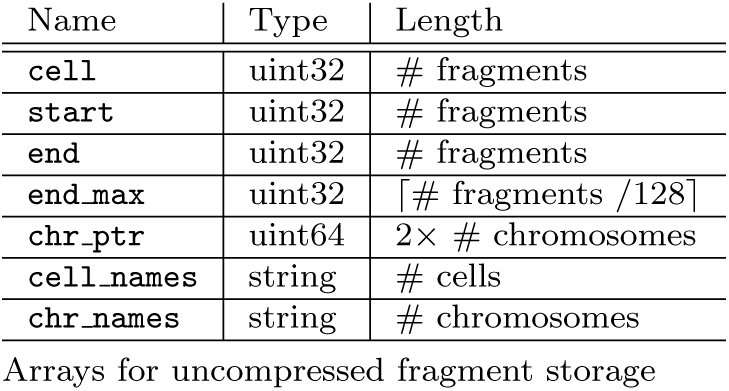

These arrays have the following contents:

- cell: List of numeric cell IDs, one per fragment. The smallest cell ID is 0.
- start: List of fragment start coordinates. The first base in a chromosome is 0.
- end: List of fragment end coordinates. The base of the end coordinate is one past the last base in the fragment.
- end max: end max[i] is the maximum end coordinate of all fragments from the start of the chromosome to the fragment at index i*128-127. If multiple chromosomes have fragments in a given chunk of 128 fragments, end max is the maximum of all those end coordinates. The end max array allows for quickly seeking to fragments overlapping a given genomic region.
- chr ptr: chr ptr[2*i] is the index of the first fragment in chromosome i in the cell, start, and end arrays. chr ptr[2*i + 1]-1 is the index of the last fragment in chromosome i. Fragments need not necessarily be sorted in order of increasing chromosome ID, though all fragments for a given chromosome must still be stored contiguously. This allows logically re-ordering chromosomes at write-time even if the input data source does not support reading chromosomes out-of-order (i.e. 10x fragment files without a genome index).
- cell names: string identifiers for each numeric cell ID.
- chr names: string identifiers for each numeric chromosome ID.

Compressed fragments are stored with the following modifications:

- cell is replaced with cell data, cell idx, and cell idx offsets, compressed according to BP-128 encoding.
- start is replaced with start data, start idx, start idx offsets, and start starts, compressed according to BP-128d1 encoding.
- end is replaced with end data, end idx, and end idx offsets, which stores start - end for each fragment, encoded using BP-128 encoding.

The current version string is equal to unpacked-fragments-v2 for uncompressed fragments, and packed-fragments-v2 for compressed fragments. In v1 formats, the only difference is that chr ptr had type uint32.

---

**Fig. S1.**
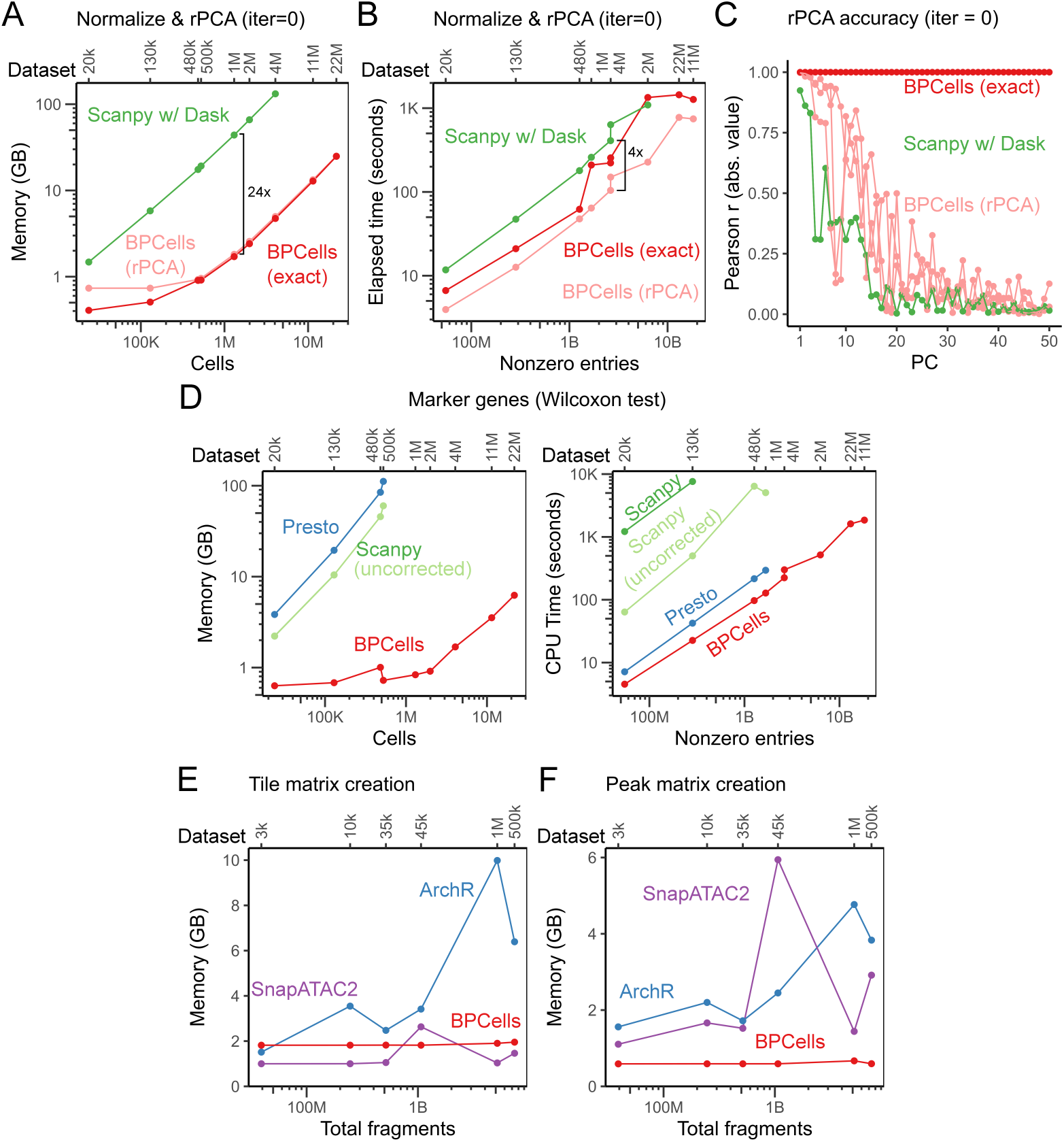
Additional disk-backed streaming performance. (a+b) BPCells compared to the Scanpy with Dask integration using the same randomized PCA algorithm with no accuracy-improving iterations, comparing (a) memory usage and (b) execution time. Note differing x-axis scales since CPU time scales more directly with nonzero entries than cells. (c) Accuracy for the 0-iteration rPCA is low compared to non-randomized solvers. (d) Performance of Wilcoxon marker gene test. Scanpy light green line shows performance when omitting the statistical correction for tied zero values. (e+f) Memory usage for ATAC-seq matrix creation routines.

**Fig. S2.**
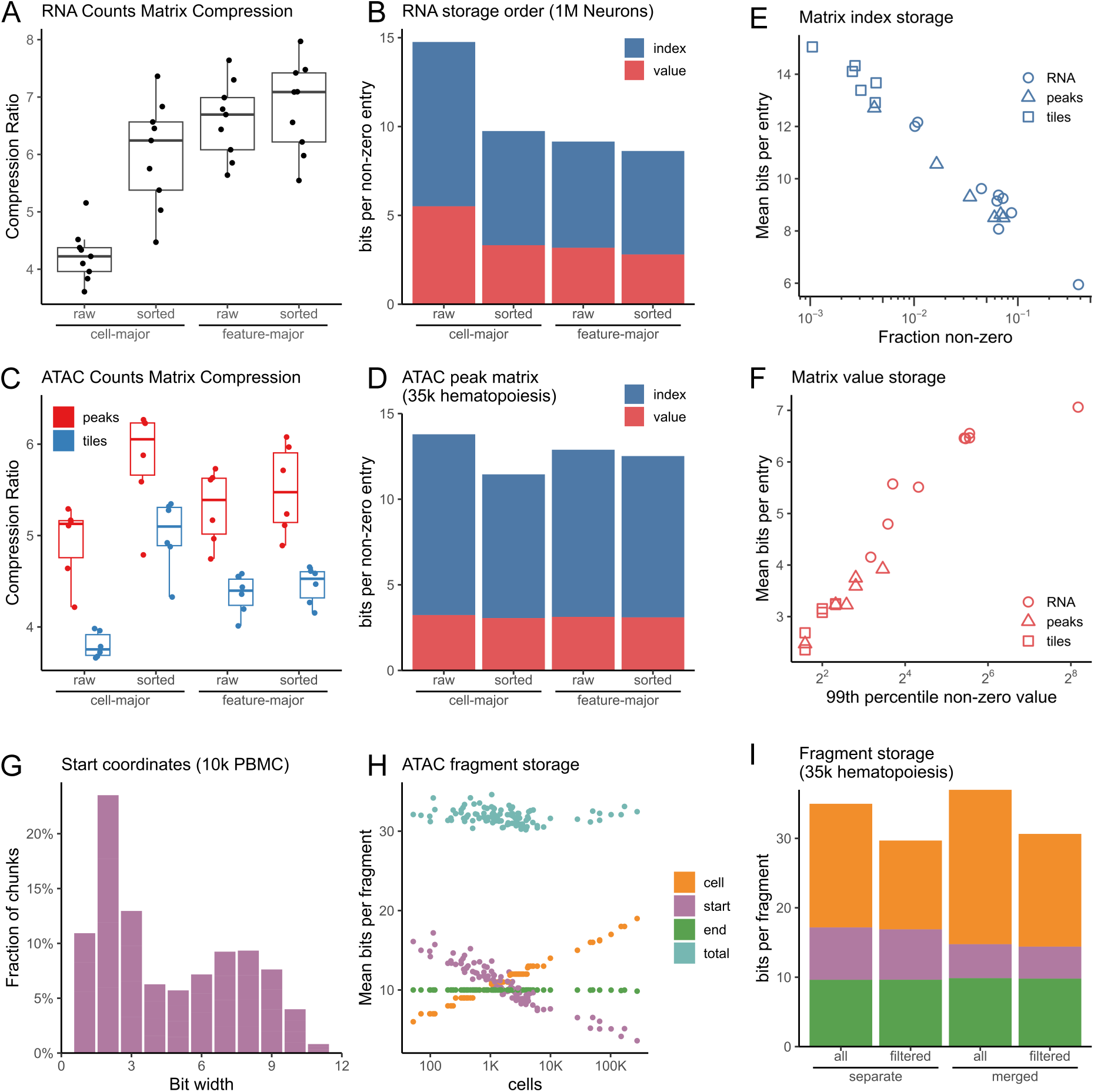
Bitpacking compression space usage properties. (a) Comparison of compression ratio across nine scRNA-seq counts matrices, comparing cell-major and feature-major storage orders with and without sorting applied to the non-major axis. (b) Average storage usage per non-zero matrix entry on the 1M neuron RNA-seq counts matrix. (c) Comparison of compression ratios across peak and tile matrices for six scATAC-seq datasets. (d) Average storage usage per non-zero matrix entry on the 35k hematopoiesis scATAC-seq peak matrix. (e) Scaling of bits to store row index for each non-zero matrix entry relative to the fraction of non-zero entries, shown across all scRNA-seq and ATAC-seq matrices. (f) Scaling of bits to store value for each non-zero matrix entry relative to the 99th percentile non-zero value, shown across all scRNA-seq and ATAC-seq matrices. (g) Distribution of bit widths used for storing start coordinate of scATAC-seq fragment data, shown across all 128-fragment chunks in the 10k PBMC dataset. (h) Storage usage for scATAC-seq fragment data across each sample in the 1M brain dataset, after filtering out non-cell barcodes. (i) Storage usage for scATAC-seq fragment data in the 35k hematopoiesis dataset when storing samples separately or merged, with and without filtering to remove non-cell barcodes.

**Fig. S3.**
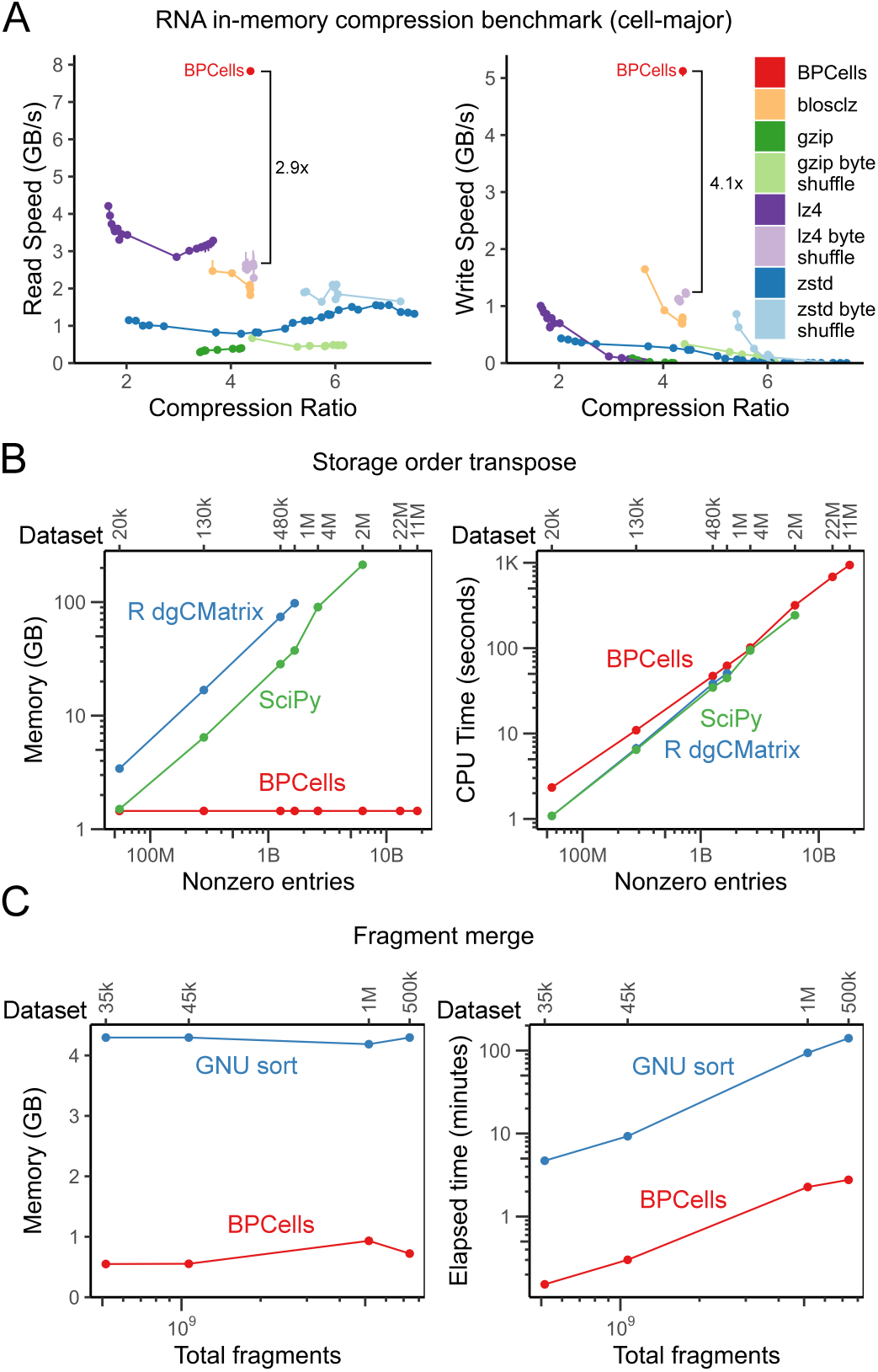
Additional compressed file performance. (a) Compression algorithm compute-efficiency benchmark on a 130k cell scRNA-seq matrix stored in cell-major order. Read/write speeds measured in uncompressed bytes, and compression ratio as uncompressed size/compressed size. (b) Storage order transposition time and memory use, compared to in-memory sparse matrix transposition. (c) Comparison of time and memory usage to merge fragment files while maintaining genome-sorted order.

**Fig. S4.**
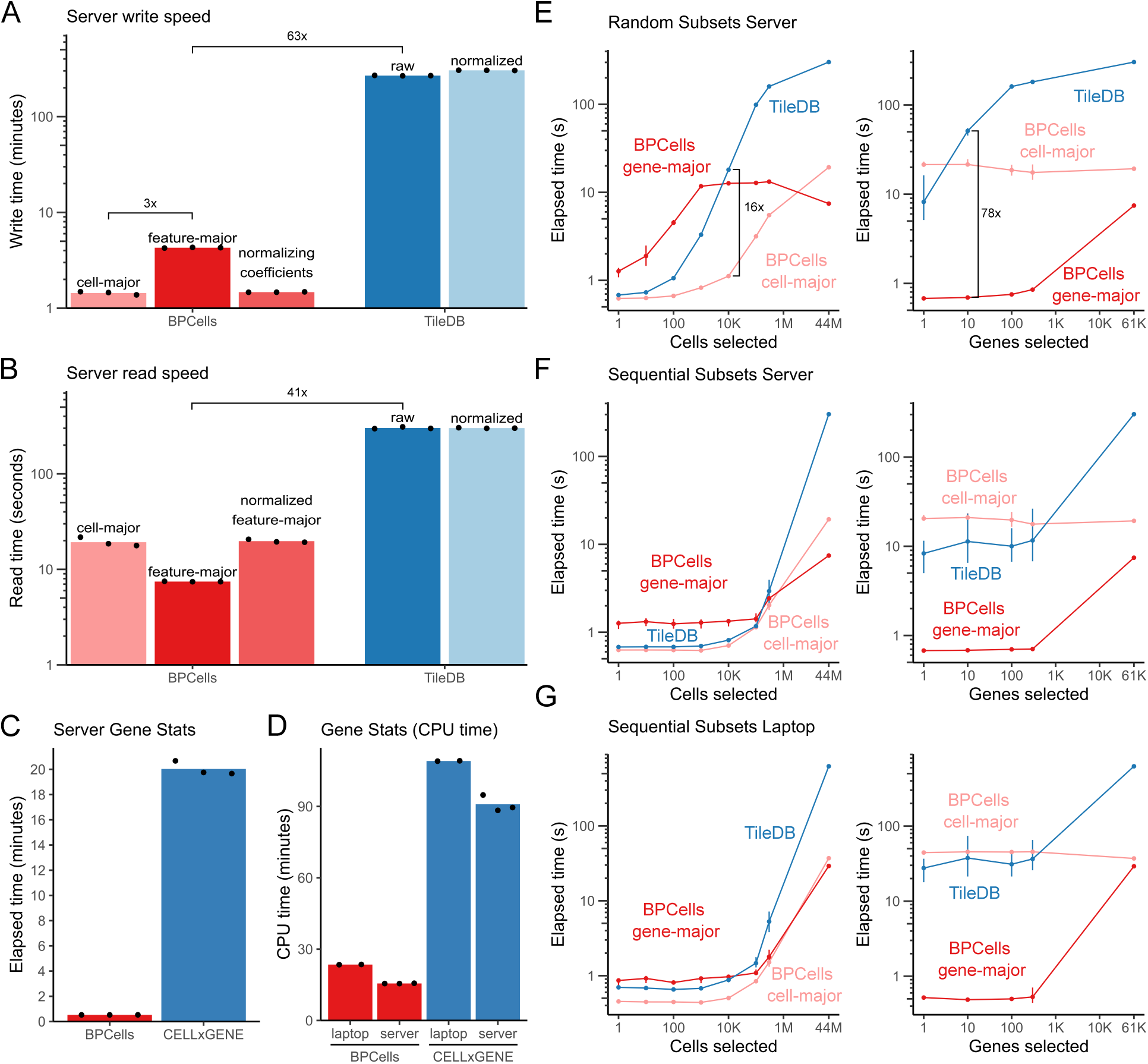
Server performance of 44M cell analysis. (a-b) Performance of BPCells sparse matrix formats and storage layouts compared to TileDB, comparing (a) write speed and (b) read speed (fully sequential) on a server. (c) Speed to calculate per-gene mean and variance on a server. (d) CPU time to calculate per-gene mean and variance on a laptop or server. (e) Read speed for random subset queries of cells and genes on a server. (f-g) Read speed for sequential subset queries of cells and genes on (f) a server and (g) a laptop.

**Table S1.** RNA benchmark datasets.

| Dataset | Download link | Cells | Genes | Median reads | Median genes | Non-zero entries | Fraction non-zero |
| --- | --- | --- | --- | --- | --- | --- | --- |
| 20k pbmc (10x Genomics) | ↗ | 23,837 | 36,601 | 6,522 | 2,020 | 54.9M | 6.3% |
| 130k thymus atlas [35] | ↗ | 129,362 | 33,694 | 5,497 | 1,830 | 284.2M | 6.5% |
| 480k tabula sapiens [36] | ↗ | 483,152 | 58,870 | 8,450 | 2,221 | 1.3B | 4.5% |
| 500k drugscreen (10x Genomics) | ↗ | 525,251 | 36,613 | 13,960 | 3,159 | 1.7B | 8.7% |
| 1M neurons (10x Genomics) | ↗ | 1,306,127 | 27,998 | 4,226 | 1,927 | 2.6B | 7.2% |
| 2M perturbseq [37] | ↗ | 1,989,578 | 8,248 | 11,304 | 3,215 | 6.3B | 38.3% |
| 4M fetal [38] | ↗ | 4,062,980 | 63,561 | 668 | 488 | 2.6B | 1.0% |
| 11M jax [39] | ↗ | 11,441,407 | 24,552 | 2,530 | 1,467 | 18.3B | 6.5% |
| 22M pansci [40] | ↗ | 21,786,931 | 55,416 | 1,034 | 460 | 13.1B | 1.1% |

**Table S2.** ATAC benchmark datasets.

| Dataset | Fragments download | Cells | Fragments | Median fragments | Total barcodes | Un-filtered fragments |
| --- | --- | --- | --- | --- | --- | --- |
| 3k pbmc (10x Genomics) | ↗ | 2,711 | 39,170,115 | 14,476 | 462,264 | 44,096,101 |
| 10k pbmc (10x Genomics) | ↗ | 10,246 | 245,922,805 | 22,218 | 549,529 | 257,551,492 |
| 35k hematopoiesis [41] | ↗ | 36,766 | 515,398,491 | 11,075 | 3,530,935 | 932,788,498 |
| 45k pancreas (ENCODE [42]) | ↗ | 45,282 | 1,065,739,414 | 15,103 | 6,291,978 | 1,563,508,448 |
| 500k heart (ENCODE [42]) | ↗ | 471,109 | 7,379,371,326 | 9,767 | 50,754,542 | 16,231,620,137 |
| 1M brain [43] | ↗ | 1,134,247 | 5,160,103,036 | 2,956 | 1,134,247 | 5,160,103,036 |

**Table S3.** BPCells matrix transformation operators.

| Operation | Order independent? | Sparsity preserving? | Sparse mat-vec product? | Notes |
| --- | --- | --- | --- | --- |
| $x \times c, x \div c$ | ✓ | ✓ | ✓ | $c$ is per-row or per-column constant |
| $x + c, x - c$ | ✓ | ✗ | ✓ | $c$ is per-row or per-column constant |
| $\log(x + 1), \exp(x) - 1$ | ✓ | ✓ | ✓ | |
| $x^c$ | ✓ | ✓ | ✓ | $c > 0$ is a user-supplied constant |
| Round, binarize | ✓ | ✓ | ✓ |  |
| $\min(x, c)$ | ✓ | ✓ | ✓ | $c > 0$ is a per-row or per-column constant |
| SCTransform [33] | ✓ | ✗ | ✗ | Pearson residuals |
| Linear regression residual | ✓ | ✗ | ✓ |  |
| Sparsity-aware rank transform | ✗ | ✓ | ✓ | Offset s.t. $\text{rank}(0) - \text{offset} = 0$ |
| Sparse matrix-matrix multiply | ✗ | ✗ | ✓ | Requires matching storage order |
| Concatenate along row/column | ✗ | ✓ | ✓ | Requires matching storage order |
| Subset rows/columns | ✓ | ✓ | ✓ |  |
| Convert integer ↔ float | ✓ | ✓ | ✓ | |

**Table S4.** BPCells matrix-consuming operations.

| Operation | Order<br>independent? | Single<br>pass? | Notes |
| --- | --- | --- | --- |
| Mean, variance, non-zero counts | ✓ | ✓ | Calculate for rows and/or columns |
| Vector, dense matrix multiply | ✓ | ✓ | Left or right multiply |
| PCA, singular value decomposition | ✓ | ✗ |  |
| Maximum per row/column | ✓ | ✓ |  |
| Quantile per row/column | ✗ | ✓ |  |
| Wilcoxon rank sum test | ✗ | ✓ | Requires feature-major order |
| User-supplied row/col reduction | ✗ | ✓ | Arbitrary operations at reduced speed |
| Pseudobulk sum, mean, variance | ✓ | ✓ |  |
| Save to disk (same ordering) | ✓ | ✓ |  |
| Save to disk (change ordering) | ✓ | ✗ |  |

**Table S5.**
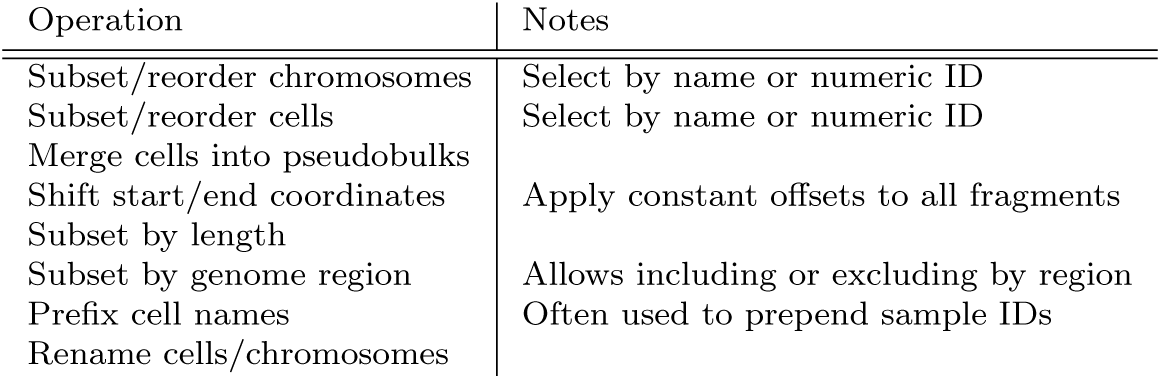
BPCells fragment transformation operators.

| Operation | Notes |
| --- | --- |
| Subset/reorder chromosomes | Select by name or numeric ID |
| Subset/reorder cells | Select by name or numeric ID |
| Merge cells into pseudobulks |  |
| Shift start/end coordinates | Apply constant offsets to all fragments |
| Subset by length |  |
| Subset by genome region | Allows including or excluding by region |
| Prefix cell names | Often used to prepend sample IDs |
| Rename cells/chromosomes |  |

**Table S6.** BPCells fragment-consuming operations.

| Operation | Output type | Notes |
| --- | --- | --- |
| Peak / tile matrix | streaming matrix | Requires BPCells-format inputs |
| Peak calling | data frame |  |
| Pseudobulk coverage tracks | bedgraph file | Coverage of Tn5 insertions |
| Pseudobulk coverage plots | plot | Calculates high-resolution tile matrix over target region |
| Gene activity matrix | streaming matrix | Matches ArchR methodology [12] |
| Cell QC stats | data frame | TSS enrichment, nucleosomal proportions, etc. matching ArchR [12] |
| Fragment length histogram | numeric vector |  |
| TF or TSS footprinting | data frame | Per-group footprinting around user-supplied genomic coordinates |
| Save to disk | fragment file | Multiple formats supported |

**Table S7.**
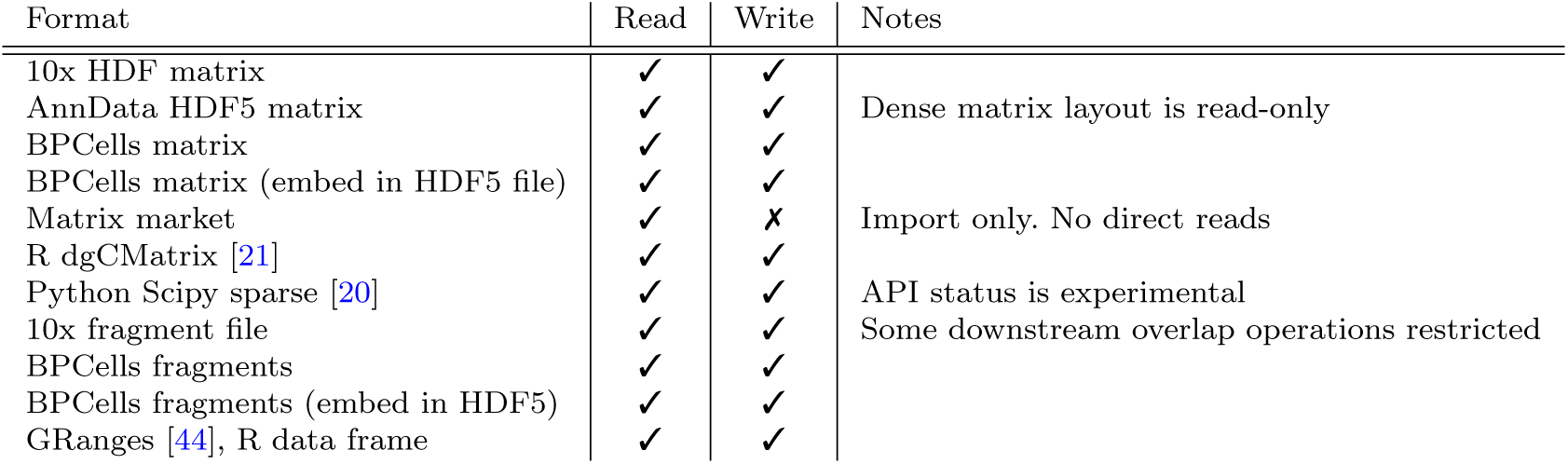
BPCells supported file formats.

## References

[1] Chan Zuckerberg Initiative. CZ CELLxGENE Discover (2023). https://cellxgene.cziscience.com/.

[2] Butler, A., Hoffman, P., Smibert, P., Papalexi, E. & Satija, R. Integrating single-cell transcriptomic data across different conditions, technologies, and species. Nature Biotechnology 36, 411–420 (2018). https://www.nature.com/articles/nbt.4096.

[3] Wolf, F. A., Angerer, P. & Theis, F. J. SCANPY: Large-scale single-cell gene expression data analysis. Genome Biology 19, 15 (2018). 10.1186/s13059-017-1382-0.

[4] Clark, I. C. et al. Microfluidics-free single-cell genomics with templated emulsification. Nature Biotechnology 41, 1557–1566 (2023). https://www.nature.com/articles/s41587-023-01685-z.

[5] Martin, B. K. et al. Optimized single-nucleus transcriptional profiling by combinatorial indexing. Nature Protocols 18, 188–207 (2023). https://www.nature.com/articles/s41596-022-00752-0.

[6] Lemire, D. & Boytsov, L. Decoding billions of integers per second through vectorization. Software: Practice and Experience 45, 1–29 (2015). https://onlinelibrary.wiley.com/doi/abs/10.1002/spe.2203.

[7] Pag’es, H., Hickey, P. & Lun, A. DelayedArray: A unified framework for working transparently with on-disk and in-memory array-like datasets. Bioconductor version: Release (3.16) (2023). https://bioconductor.org/packages/DelayedArray/.

[8] McKinney, W. Wes McKinney - Apache Arrow and the “10 Things I Hate About pandas” (2017). https://wesmckinney.com/blog/apache-arrow-pandas-internals/.

[9] Rocklin, M. Dask: Parallel Computation with Blocked algorithms and Task Scheduling, 126–132 (Austin, Texas, 2015). https://conference.scipy.org/proceedings/scipy2015/matthewrocklin.html.

[10] Gold, I. & Virshup, I. Using dask with Scanpy (2024). https://web.archive.org/web/20250117053910/ https://scanpy.readthedocs.io/en/stable/tutorials/experimental/dask.html.

[11] Halko, N., Martinsson, P.-G. & Tropp, J. A. Finding structure with randomness: Probabilistic algorithms for constructing approximate matrix decompositions (2009). https://arxiv.org/abs/0909.4061v2.

[12] Granja, J. M. et al. ArchR is a scalable software package for integrative single-cell chromatin accessibility analysis. Nature Genetics 53, 403–411 (2021). https://www.nature.com/articles/s41588-021-00790-6.

[13] Zhang, K., Zemke, N. R., Armand, E. J. & Ren, B. A fast, scalable and versatile tool for analysis of single-cell omics data. Nature Methods 21, 217–227 (2024). https://www.nature.com/articles/s41592-023-02139-9.

14. Korsunsky, I., Nathan, A., Millard, N. & Raychaudhuri, S. Presto scales Wilcoxon and auROC analyses to millions of observations (2019). https://www.biorxiv.org/content/10.1101/653253v1.

[15] Deutsch, L. P. GZIP file format specification version 4.3. Request for Comments RFC 1952, Internet Engineering Task Force (1996). https://datatracker.ietf.org/doc/rfc1952.

[16] Buenrostro, J. D., Giresi, P. G., Zaba, L. C., Chang, H. Y. & Greenleaf, W. J. Transposition of native chromatin for fast and sensitive epigenomic profiling of open chromatin, DNA-binding proteins and nucleosome position. Nature Methods 10, 1213–1218 (2013). https://www.nature.com/articles/nmeth.2688.

[17] Collet, Y. RealTime Data Compression: LZ4 explained (2011). http://fastcompression.blogspot.com/2011/05/lz4-explained.html.

[18] Collet, Y. Zstandard - Real-time data compression algorithm (2016). https://facebook.github.io/zstd/.

[19] Alted, F. Why Modern CPUs Are Starving and What Can Be Done about It. Computing in Science & Engineering 12, 68–71 (2010).

[20] Virtanen, P. et al. SciPy 1.0: Fundamental algorithms for scientific computing in python. Nature Methods 17, 261–272 (2020). 10.1038/s41592-019-0686-2.

[21] Bates, D., Maechler, M. & Jagan, M. Matrix: Sparse and Dense Matrix Classes and Methods (2024). https://cran.r-project.org/web/packages/Matrix/index.html.

[22] Haertel, M. & Eggert, P. GNU sort (2022). https://www.gnu.org/software/coreutils/.

[23] Wolen, A., Eddelbuettel, D., Hoffman, P. & Kerl, J. Tiledbsoma: –TileDB’ Stack of Matrices, Annotated (’SOMA*’)* (2024). https://github.com/single-cell-data/TileDB-SOMA.

[24] Papadopoulos, S., Datta, K., Madden, S. & Mattson, T. The TileDB array data storage manager. Proc. VLDB Endow. 10, 349–360 (2016). https://dl.acm.org/doi/10.14778/3025111.3025117.

[25] Hao, Y. et al. Dictionary learning for integrative, multimodal and scalable single-cell analysis. Nature Biotechnology 42, 293–304 (2024). https://www.nature.com/articles/s41587-023-01767-y.

[26] Lee, D. & Seung, H. S. Algorithms for non-negative matrix factorization. Advances in neural information processing systems 13 (2000).

[27] Stuart, T., Srivastava, A., Madad, S., Lareau, C. A. & Satija, R. Single-cell chromatin state analysis with Signac. Nature Methods 18, 1333– 1341 (2021). https://www.nature.com/articles/s41592-021-01282-5.

[28] Friedman, J., Hastie, T. & Tibshirani, R. Regularization paths for generalized linear models via coordinate descent. Journal of statistical software 33, 1 (2010).

[29] Hie, B., Cho, H., DeMeo, B., Bryson, B. & Berger, B. Geometric Sketching Compactly Summarizes the Single-Cell Transcriptomic Landscape. Cell Systems 8, 483–493.e7 (2019). https://www.sciencedirect.com/science/article/pii/S2405471219301528.

[30] Lopez, R., Regier, J., Cole, M. B., Jordan, M. I. & Yosef, N. Deep generative modeling for single-cell transcriptomics. Nature Methods 15, 1053–1058 (2018). https://www.nature.com/articles/s41592-018-0229-2.

[31] Skibinski, P. Inikep/lzbench (2025). https://github.com/inikep/lzbench.

[32] Wassenberg, J. Highway. Google (2025). https://github.com/google/highway.

[33] Hafemeister, C. & Satija, R. Normalization and variance stabilization of single-cell RNA-seq data using regularized negative binomial regression. Genome Biology 20, 296 (2019). 10.1186/s13059-019-1874-1.

[34] Ammon, P. Ridiculousfish/libdivide (2011). https://github.com/ridiculousfish/libdivide.

[35] Park, J.-E., et al. A cell atlas of human thymic development defines T cell repertoire formation. Science 367, eaay3224 (2020). https://www.science.org/doi/10.1126/science.aay3224.

[36] THE TABULA SAPIENS CONSORTIUM. The Tabula Sapiens: A multiple-organ, single-cell transcriptomic atlas of humans. Science 376, eabl4896 (2022). https://www.science.org/doi/10.1126/science.abl4896.

[37] Replogle, J. M. et al. Mapping information-rich genotype-phenotype landscapes with genome-scale Perturb-seq. Cell 185, 2559–2575.e28 (2022). https://www.sciencedirect.com/science/article/pii/S0092867422005979.

[38] Cao, J., et al. A human cell atlas of fetal gene expression. Science 370, eaba7721 (2020). https://www.science.org/doi/10.1126/science.aba7721.

[39] Qiu, C. et al. A single-cell time-lapse of mouse prenatal development from gastrula to birth. Nature 626, 1084–1093 (2024). https://www.nature.com/articles/s41586-024-07069-w.

[40] Zhang, Z., et al. A panoramic view of cell population dynamics in mammalian aging. Science 387, eadn3949 (2025). https://www.science.org/doi/abs/10.1126/science.adn3949.

[41] Granja, J. M. et al. Single-cell multiomic analysis identifies regulatory programs in mixed-phenotype acute leukemia. Nature Biotechnology 37, 1458–1465 (2019). https://www.nature.com/articles/s41587-019-0332-7.

[42] Moore, J. E. et al. Expanded encyclopaedias of DNA elements in the human and mouse genomes. Nature 583, 699–710 (2020). https://www.nature.com/articles/s41586-020-2493-4.

[43] Li, Y. E., et al. A comparative atlas of single-cell chromatin accessibility in the human brain. Science 382, eadf7044 (2023). https://www.science.org/doi/10.1126/science.adf7044.

[44] Lawrence, M., et al. Software for Computing and Annotating Genomic Ranges. PLOS Computational Biology 9, e1003118 (2013). https://journals.plos.org/ploscompbiol/article?id=10.1371/journal.pcbi.1003118.

